# TARGETING THE CAPSULE OF *KLEBSIELLA PNEUMONIAE* WITH A CATIONIC CR3-BINDING PROTEIN ENHANCES PHAGOCYTOSIS AND PROMOTES BACTERIAL CLEARANCE AND SURVIVAL IN A MOUSE SEPSIS MODEL

**DOI:** 10.64898/2026.07.14.737852

**Authors:** Nataly P. Podolnikova, Iryna Klymenko, James Alagna, Cyrus Diercksmeier, Arnat Balabiyev, David Richardson, Tatiana P. Ugarova

**Affiliations:** From the School of Life Sciences, Arizona State University, Tempe, AZ 85287, United States; bioSyntagma Inc., Tempe, AZ 85282, United States

**Keywords:** *Klebsiella pneumoniae*, CR3, Mac-1, phagocytosis, antibiotic resistance, PF4, AMP

## Abstract

Platelet Factor 4 (PF4), a cationic antimicrobial peptide, serves as a ligand for the myeloid-specific phagocytic receptor CR3 (Mac-1, CD11b/CD18). We previously demonstrated that recombinant dimeric PF4 (rdPF4) functions as a bacterial opsonin, enhancing phagocytosis of Gram-positive *Staphylococcus aureus* and facilitating clearance of both antibiotic-susceptible and methicillin-resistant *S. aureus* in a mouse model of infectious peritonitis. In this study, we examined whether rdPF4 is pathogen-agnostic by assessing its effect on phagocytosis of Gram-negative encapsulated *Klebsiella pneumoniae*, a WHO Bacterial Priority Pathogen. We demonstrate that rdPF4 enhances CR3-mediated phagocytosis of both live and heat-inactivated high-virulence K2 and low-virulence K3 strains of *K. pneumoniae* by various mouse and human macrophage cell lines, as well as primary neutrophils and macrophages. It also increased phagocytosis of carbapenem-resistant *K. pneumoniae*. rdPF4 did not directly kill bacteria but acted as an opsonin binding to the negatively charged bacterial capsule and creating recognition sites for CR3 on leukocytes. In a mouse sepsis model, a single dose of rdPF4 significantly enhanced bacterial clearance from the lungs, liver, and peritoneum and reduced bacteremia. Histological analyses showed that rdPF4 provided substantial protection to lung and liver tissues against *K. pneumoniae*-induced damage. Consistent with these findings, rdPF4 treatment increased the survival rates of infected mice. These results show that rdPF4 effectively targets the capsule, a key virulence factor of *K. pneumoniae*, thereby reducing the bacterium’s ability to evade the host immune response. Overall, the data suggest a common mechanism in which cationic rdPF4, by binding to the negatively charged surfaces of both Gram-negative and Gram-positive bacteria, diminishes their antiphagocytic properties.

## INTRODUCTION

The Gram-negative, encapsulated bacterium *Klebsiella pneumoniae* is a significant cause of human disease. Although considered an opportunistic pathogen that colonizes mucosal surfaces in healthy individuals, it can disseminate to other tissues, leading to life-threatening infections such as pneumonia, bloodstream and urinary tract infections, meningitis, endophthalmitis, and sepsis (1–3). Additionally, *K. pneumoniae*-induced liver abscess is among the most severe complications caused by this pathogen (4). Infections caused by *K. pneumoniae* are often difficult to treat because these organisms are frequently resistant to multiple antibiotics (3, 5).

It is well documented that the bacterial capsule, especially prominent in *K. pneumoniae*, is a key virulence factor that enables bacteria to evade host defenses (1, 3). It has been suggested that the capsule, composed of negatively charged capsular polysaccharides (CPS), can resist phagocytosis by repelling the negatively charged surfaces of phagocytes, with more highly charged CPS being more effective (reviewed in (6)). Other mechanisms include limiting complement-mediated killing by sterically hindering the membrane attack complex (MAC) from reaching the outer membrane (3, 7), and reducing opsonophagocytosis by masking C3b/iC3b and capsule-specific antibodies beneath the capsule layer, thereby preventing their recognition by phagocytic complement receptors 1 and 3 (CR1 and CR3) (3, 8). Numerous studies using mouse models have shown that encapsulated *K. pneumoniae* strains have higher bacterial loads and mortality rates compared to less virulent, acapsular mutants (9–13). Moreover, the link between CPS production and increased pathogenicity of *K. pneumoniae* has been reported in some studies (14, 15) but not in others (16). Specifically, high-virulence strains, such as the K1 and K2 serotypes, which cause about 80% of severe infections (17–19), produce a thick, viscous polysaccharide coat (hypercapsule) on their surface. The presence of the hypercapsule has been associated with greater resistance to phagocytosis by human neutrophils and macrophages compared to classical strains (20–22). The role of *K. pneumoniae’s* capsular components in virulence has made them promising targets for therapeutics and vaccines. In particular, several studies have demonstrated that anti-CPS antibodies can protect experimental animals from *K. pneumoniae* infection (5).

Along with the widely accepted proposal that the capsule limits phagocytosis by repelling the negatively charged surfaces of phagocytic leukocytes, the unique ligand-recognition specificity of the major phagocytic receptor, CR3 (also known as Mac-1, integrin αMβ2, CD11b/CD18), may help explain how bacteria can evade these cells. CR3, which is abundantly expressed on the surface of myeloid leukocytes, such as neutrophils and monocytes/macrophages, is a multiligand receptor that can bind its ligands through two distinct mechanisms. In the first, it binds several ligands, specifically iC3b, a fragment of C3b complement, by coordinating a glutamic acid in its thioester domain through MIDAS (a metal-dependent adhesion site) located in the ligand-binding αMI domain of CR3 (23, 24). However, the αMI domain can also bind “noncomplement” ligands. In these ligands, the αMI-domain binds sequences enriched in basic and hydrophobic residues while strongly disfavoring negatively charged residues (25). It has been shown that the isolated αMI domain and leukocytes that express CR3 generally do not adhere to surfaces coated with negatively charged polysaccharides(26–28). In particular, the recombinant αMI domain does not bind to carboxymethylated dextran, and macrophages and CR3-expressing HEK293 cells do not adhere to surfaces coated with heparan sulfate and aggrecan. Consequently, the capsule can resist phagocytosis not only by repelling the negatively charged glycocalyx on phagocytes but also by preventing CR3 from binding to the negatively charged bacterial surface.

Among the noncomplement ligands that contain multiple positively charged CR3 recognition sequences are cationic proteins and peptides that, under physiological conditions, are stored within leukocyte and platelet granules and released during the inflammatory response. These molecules include leukocyte myeloperoxidase, neutrophil elastase, cathepsin G, azurocidin, LL-37, human β-defensisn, proteinase 3, dynorphin, and others (29, 30). We recently identified platelet protein PF4 (platelet factor 4), a small (7.8 kDa) cationic protein present in mammalian platelet α-granules at unusually high concentrations, as a CR3 ligand and demonstrated that, in addition to mediating various leukocyte responses, it acts as a potent opsonin (28, 30). PF4 possesses two sequences, each containing a core of positively charged residues flanked by hydrophobic residues, thereby conforming to the αMI domain’s recognition specificity for noncomplement ligands. Because of their cationic nature, these sequences can also bind to the bacterial surface. Therefore, having two segments that serve as interchangeable binding sites for CR3 and bacteria makes PF4 an excellent opsonin, marking bacteria and making them attractive targets for phagocytes. Indeed, we recently showed that recombinant dimeric PF4 (rdPF4) enhanced phagocytosis of both nonencapsulated and encapsulated *Staphylococcus aureus* by CR3-expressing leukocytes, and significantly increased the clearance of antibiotic-susceptible bacteria, as well as its methicillin-resistant (MRSA) strain, markedly improving survival rates in the murine peritonitis model (28).

In this study, we postulated that targeting the negatively charged bacterial capsule of Gram-negative bacteria with rdPF4 could also overcome its antiphagocytic properties. To investigate this, we examined the effect of rdPF4 on the phagocytosis of several encapsulated *K. pneumoniae* strains, including K2 and K3 serotypes, by various CR3-expressing leukocytes *in vitro*. Additionally, we evaluated the role of rdPF4 as an enhancer of bacterial clearance in a mouse sepsis model, its ability to protect organs from injury caused by *K. pneumoniae* infection, and its potential to improve survival.

## Materials and methods

### Reagents

The rat mAb M1/70 and mouse mAb 44a, which recognize the mouse and human αM (CD11b) subunit of CR3 (integrin Mac-1), respectively, were purified from conditioned media of hybridoma cells obtained from the American Type Culture Collection (ATCC, Manassas, VA) using protein A agarose. The Alexa Fluor 488-conjugated anti-Ly6G mouse mAb (catalog #127625), directed against neutrophil-specific lymphocyte antigen 6, locus G, and Ultra-LEAF purified anti-mouse Ly6G (clone 1A8, catalog #127632) were from BioLegend (San Diego, CA). The secondary antibodies, Alexa Fluor 488-conjugated goat anti-mouse IgG (H+L) (catalog **#**A-10680), Alexa Fluor 633-conjugated goat anti-rabbit IgG (catalog #A-21071), and Alexa Fluor 633-conjugated goat anti-rat IgG (catalog #A-21094) were from Thermo Fisher. The polyclonal anti-PF4 antibody was raised in rabbits using recombinant PF4 as an antigen. NHS-Fluorescein (catalog #46410), Alexa Fluor 568-conjugated phalloidin (catalog #A12380), and pHrodo iFL Red STP ester (catalog #P36011) were from Thermo Fisher. Zwittergent 3-14 (catalog #693017), 3-phenylphenol (catalog #262250), glucuronic acid (catalog #G5269), heparan sulfate sodium salt (catalog #7640), and 4% Brewer thioglycolate (TG) solution were from Sigma-Aldrich (St. Louis, MO). The macrophage Depletion Kit (Clodrosome) was from Encapsula Nano Science (catalog # CLD-8901).

### Bacterial strains and growth conditions

Three different strains of *K. pneumoniae* were obtained from ATCC: low-virulence K3 serotype of *K. pneumoniae* (lv*Kp*) subsp. pneumoniae (Schroeter) Trevisan (ATCC 13883), high-virulence K2 serotype of *K. pneumoniae* (hv*Kp*) subsp. pneumoniae (Schroeter) Trevisan (ATCC 43816), and carbapenem-resistant *K. pneumoniae* (Schroeter) Trevisan (catalog #BAA-1705). Bacteria were grown overnight at 37 °C on Luria-Bertani (LB) agar plates. A single colony was transferred to LB medium and incubated with shaking in screw-cap 15-mL plastic tubes containing 3 mL of medium for 16 hours at 37 °C. The culture was diluted 100-fold in fresh media and grown at 37 °C until reaching mid-log phase. The bacteria were harvested by centrifugation at 6,000×g for 5 minutes, washed with PBS, and adjusted to the desired CFU/mL. Colony-forming units (CFU) per milliliter were calculated by counting colonies from serial dilutions on agar plates incubated overnight at 37 °C.

### Mice

C57BL/6 mice were obtained from The Jackson Laboratory (Bar Harbor, ME) or bred at Arizona State University animal facility with mating pairs originally purchased from The Jackson Laboratory. All experiments were conducted using male and female mice aged 8 to 12 weeks, with age- and sex-matched groups selected for side-by-side comparison. The animals were housed in the Animal Facility at Arizona State University, where they were kept under controlled temperature (22 °C) and humidity conditions on a 12-hour light/dark cycle. This study was conducted in strict accordance with the recommendations of the Guide for the Care and Use of Laboratory Animals of the National Institutes of Health. The use of mice was approved by the Institutional Animal Care and Use Committee of Arizona State University (#23-1964R and #26-2176R). Mice were consistently monitored for signs of distress throughout the experiments and were removed from the experiment and euthanized by overdose of isoflurane inhalation (5%) to prevent unnecessary suffering, in accordance with the approved institutional animal care protocol.

To generate neutrophil-depleted mice, animals were injected with 100 µg of anti-Ly6G antibody (clone 1A8) 24 hours before infection with *K. pneumoniae*. Depletion was assessed by measuring neutrophil counts in peritoneal lavage after bacterial infection. Macrophages were depleted using the Clodrosome macrophage depletion kit. Mice received a retro-orbital injection of 0.2 mL of either control liposomes or Clodronate-containing liposomes, followed 24 hours later by an intraperitoneal injection. *K. pneumoniae* was administered on day 3 after the initial liposome treatment.

### Cells

The IC-21 murine macrophage cell line, human monocytic U937 and THP-1 cells, and A549 lung epithelial cells were obtained from ATCC and cultured in RPMI supplemented with 10% FBS and antibiotics. Wild-type human embryonic kidney (WT-HEK293) cells and HEK293 cells stably expressing CR3 (CR3-HEK293) (31) were grown in DMEM/F12 with 10% FBS and antibiotics. Resident peritoneal macrophages were obtained from mice by lavage with cold PBS containing 5 mM EDTA as described (32). Peritoneal lavage containing neutrophils and monocytes/macrophages was isolated from mice 4 hours and 3 days, respectively, after intraperitoneal injection of 0.5 mL of a 4% thioglycollate (TG) solution. The cell population in the 4-hour lavage included both Ly6G^+^CD11b^+^ neutrophils and Ly6G^-^CD11b^+^ monocyte/macrophages (28). Human neutrophils, PBMCs, and red blood cells were isolated from peripheral blood drawn from healthy volunteers under sterile conditions, as described (33).

### Preparation of recombinant PF4

Recombinant PF4 was expressed and purified according to established procedures (28). Briefly, the open reading frame of human PF4 cDNA (E1-S70) was cloned into the pET-15b vector (Novagen, Madison, WI) and transformed into Origami B (DE3) competent cells (Novagen, Madison, WI). The protein was induced with 0.5 mM IPTG, and rPF4 was purified from soluble cell lysate fractions by affinity chromatography using a 5-mL HiTrap Heparin-agarose column (GE Healthcare). The protein’s purity was verified by electrophoresis and Western blotting with a polyclonal antibody against human PF4. Size-exclusion chromatography on a BioSec-3 column revealed that the recombinant protein consists of ∼80% dimers and 20% monomers; it is therefore referred to as rdPF4. Endotoxin-free rdPF4 was prepared using Pierce™ High-Capacity Endotoxin Removal Spin Columns according to the manufacturer’s protocol.

### Detection of rdPF4 on the surface of K. pneumoniae and mammalian cells

*K. pneumoniae* (1 × 10^8^/mL) and various human and mouse cells in PBS were incubated with rdPF4 for 30 minutes at 22 °C, then washed twice with PBS. The cells were subsequently incubated with rabbit polyclonal anti-PF4 antibody (1:250) for 30 minutes at 22°C, followed by incubation with Alexa Fluor 488-conjugated secondary antibody. After washing with PBS, the cells were resuspended in 1% paraformaldehyde and then deposited onto glass coverslips by cytospin at 2000×g for 5 minutes. Confocal images were captured with a Leica SP8 Confocal System (Exton, PA) using a 60x/1.4 oil objective.

### Preparation of K. pneumoniae capsular polysaccharides

Capsular polysaccharides were isolated from a low virulence strain of *K. pneumoniae* as described (34). Briefly, the capsule was extracted from the bacterial culture using Zwittergent 3-14 detergent and precipitated with ethanol. Residual proteins were removed by digestion with pronase and trypsin, followed by an additional ethanol precipitation step. The precipitate was dissolved in water and stored at −20 °C.

### Labeling of bacteria with fluorescein and pHrodo dyes

Heat-inactivated for 1 hour at 70 °C *K. pneumoniae* were washed in 0.1 M sodium bicarbonate buffer, pH 9.0, and labeled with 10 μg/mL NHS-fluorescein for 1 hour at 22 °C. The bacteria were washed to remove unbound fluorochrome, then aliquoted and stored at −20 °C until further use. In selected experiments, live *K. pneumoniae* were labeled with a pHrodo dye according to the manufacturer’s protocols, then aliquoted, and stored at −20 °C.

### Phagocytosis assays

Phagocytosis assays with fluorescein-labeled heat-inactivated *K. pneumoniae* and various CR3-expressing cells were performed using both suspended and adherent cells. In the first format, *K. pneumoniae* (10^8^ CFU) was incubated with different concentrations of rdPF4 for 1 hour at 37 °C. Aliquots (100 μL; 3×10^5^) of various cells, including IC-21 mouse macrophages, mouse peritoneal resident and inflammatory macrophages, and mouse and human neutrophils, suspended in Hank’s balanced salt solution (HBSS), were incubated with rdPF4-treated bacterial particles for 30 min at 37 °C. Ice-cold PBS (400 µL) was added to the cell mixtures to stop the phagocytosis. 50 μg/mL ethidium bromide was added to quench extracellular bacteria, and samples were analyzed using an Attune NxT flow cytometer (ThermoFisher).

For phagocytosis assays with adherent cells, IC-21 macrophages, inflammatory peritoneal mouse macrophages, and a mixture of neutrophils and monocytes in DMEM+10% FBS were allowed to adhere to acid-cleaned glass at a density of 2.5×10^5^/well for 3-5 hours. Fluorescein-labeled bacteria were incubated with 50 µg/mL rdPF4 for 1 hour at 37 °C, and then 0.5 mL aliquots were added to the cells for an additional 1 hour at 37 °C. Cells were washed three times with 1 mL of PBS, fixed with 2% paraformaldehyde in PBS for 30 min, and stained with 15 nM Alexa Fluor 568-conjugated phalloidin for 30 min at 22 °C to detect F-actin. Cells were washed twice with PBS and incubated with DAPI. ProLong Diamond Antifade Mountain was used to mount the cells on a glass slide (Thermo Scientific, Waltham, MA). Images were acquired using a Leica SP8 confocal microscope with a 60x/1.4 oil immersion objective. Phagocytosed bacteria were counted using ImageJ software.

Phagocytosis assays with heat-inactivated and live pHrodo-conjugated *K. pneumoniae* were performed using IC-21, PMA-activated THP-1 cells, TG-elicited mouse inflammatory macrophages, and a mixture of mouse neutrophils and monocytes suspended in culture medium. pHrodo-conjugated *K. pneumoniae* was incubated with different concentrations of rdPF4 for 1 hour at 37 °C. Cell aliquots (100 μL; 3×10^5^) were incubated with rdPF4-treated bacteria for 30 min at 37 °C. Ice-cold PBS (400 µL) was added to stop the reaction, and the samples were analyzed by flow cytometry.

### Adhesion assays

96-well microtiter plates (Immulon 4HBX, Thermo Labsystems, Franklin, MA) were coated with varying concentrations of capsular polysaccharide (CPS) or heparan sulfate, or CPS followed by rdPF4, for 16 hours at 4 °C, then blocked with 1.0% PVP for 1 hour at 22 °C. Cells were labeled with 5 μM calcein for 30 min at 37 °C, washed twice with Hank’s balanced salt solution containing 0.1% BSA, and then added to each well. Aliquots of 5×10^4^ cells in 100 μL were added to each well and allowed to adhere for 30 min at 37 °C. Nonadherent cells were removed by two washes with PBS, and fluorescence was quantified using a fluorescence plate reader. The number of adherent cells was determined by comparing the fluorescence of 100-μL aliquots containing a known number of labeled cells. In separate experiments, wells were precoated with different concentrations of CPS, treated with rdPF4, incubated for 3 h at 37 °C, and then subjected to adhesion as above.

### Flow Cytometry

Expression of neutrophil- and macrophage-specific antigens on the surface of leukocytes isolated from the peritoneum 4 hours after TG injection was analyzed by flow cytometry. Cells were harvested, washed twice with ice-cold HBSS, and resuspended at a concentration of 3×10^6^/ml. 300,000 cells per sample were then stained with fluorophore-conjugated monoclonal antibodies specific for cell surface receptors for 30 minutes at 4 °C in the dark. After staining, cells were washed in PBS, and data were acquired on a Thermo Fisher Scientific Attune NxT instrument, with at least 10,000 events collected per sample. Data were analyzed using FlowJo (version 10) software. Receptor expression was quantified as the percentage of positive cells and/or the median fluorescence intensity (MFI) relative to isotype controls.

### Histology and Immunohistochemistry

Isolated mouse lung and liver tissues were fixed in 10% formalin for 24 hours at 22°C. After fixation, tissues were embedded in paraffin, sectioned, and stained using a standard H&E histological procedure. Sections were mounted on cover slips and imaged using the EVOS FL Auto (Thermo Scientific, Waltham, MA) wide-field microscope with a 20x objective. To examine the presence of various cells and proteins in lung tissue, paraffin blocks were sectioned, and sections were deparaffinized and dehydrated in ethanol. After incubation in PBS with 1% BSA, sections were incubated with primary mAbs against Ly6G and F4/80 for 1 hour at 22°C, followed by incubation with secondary antibodies conjugated to Alexa Fluor 647. Samples were mounted in ProLong Diamond (Thermo Scientific) and imaged with a 20x oil-immersion objective using an SP8 confocal microscope.

### A mouse sepsis model

Infections were initiated by intraperitoneal (i.p.) injection of low-virulence (1×10^8^ CFU) or high-virulence (1×10^4^ CFU) strains of *K. pneumoniae*, followed immediately by an i.p. injection of rdPF4. CFU counts for each inoculum were determined by plating serial dilutions before infection to retroactively calculate the actual infection dose. To relieve distress, mice were injected subcutaneously with buprenorphine (0.05 mg/kg) twice daily. Mice were examined every hour for the first 8 hours, then every 6 hours thereafter for signs of illness. If a mouse showed signs of moribundity, it was euthanized. Mice were sacrificed after 24 hours, and peritoneal cavities were lavaged with 5 mL of sterile, endotoxin-free PBS containing 5 mM EDTA. Lung and liver samples were collected, and 100 μL of blood was obtained via cardiac puncture. Lung and liver tissues were minced, mechanically homogenized, and filtered through a 100-μm strainer. Aliquots of peritoneal lavage fluid, blood, and tissue suspensions were serially diluted and plated on LB agar plates. After 16 hours of incubation at 37 °C, colonies on the plates were counted. Peritoneal inflammation was assessed by counting total leukocytes in the peritoneal lavage, and differential cell counts were performed using Wright-stained cytospin preparations.

### Survival studies

Mice were inoculated intraperitoneally with low-virulence (5×10^8^ CFU), high-virulence (2×10^4^ CFU), and antibiotic-resistant BAA-1750 (5×10^7^ CFU) strains of *K. pneumoniae*, with or without endotoxin-free rdPF4 (0.6-2.5 mg/kg, 12-50 µg/mouse). Animals were monitored for signs of mortality and morbidity every hour for the first 12 hours, then four times daily for 10 days. If a mouse demonstrated signs of moribundity, it was euthanized. Kaplan-Meier survival plots were generated to assess the effect of rdPF4 on survival.

### Statistical analysis

Data are presented as the mean ± standard deviation (S.D.). Statistical comparisons between the two groups were performed using a Student’s t-test. For multiple comparisons, ANOVA was followed by Tukey’s or Dunn’s post hoc test, conducted in GraphPad Prism. A p-value less than 0.05 was considered statistically significant.

## RESULTS

### rdPF4 binding to K. pneumoniae and mammalian cells

Previous studies demonstrated that both cationic proteins, intact tetrameric PF4 isolated from human platelets(30, 35) and recombinant dimeric PF4 (rdPF4) (28), can bind to negatively charged molecules on the surfaces of selected Gram-positive (*S. aureus, S. pneumoniae*, and *L. monocytogenes*) and Gram-negative (*E. coli* and *N. meningitidis*) bacteria. Since *K. pneumoniae* is encapsulated and the capsule usually increases the negative charge of the cell surface, we examined its ability to bind rdPF4. Because biotinylation, a standard method for detecting protein molecules, modifies positively charged lysines and may reduce rdPF4’s binding to negatively charged bacterial surfaces, we performed immunostaining. Using an anti-PF4 polyclonal antibody, we showed that rdPF4 binds to the surface of *K. pneumoniae* (Fig. 1A). Because mammalian cells also express negatively charged polysaccharides and can bind PF4 (36), we compared the density of rdPF4 bound to *K. pneumoniae* and to various cultured and primary human and mouse cells. The specificity of rdPF4 binding to CR3 was confirmed using recombinant CR3-expressing HEK293 cells, which bound significantly more rdPF4 than wild-type HEK293 cells (Fig. 1B). The density of rdPF4 bound to the surface of blood cells, such as RBCs, nonactivated neutrophils, and PBMCs, was lower than that to the surface of bacteria (Fig. 1, C and D). Low binding of rdPF4 was also observed in selected cultured cells, including HEK293 and A549 epithelial cells. However, CR3-expressing leukocytes, including resident peritoneal mouse macrophages, activated human neutrophils, and mouse IC-21 plus human THP-1 macrophages, bound rdPF4, albeit at lower levels than bacteria (Fig. 1, C and D). To determine whether rdPF4 preferentially binds to the bacterial surface rather than to CR3-expressing phagocytes when both cell types are present, we incubated rdPF4 with resident mouse macrophages and *K. pneumoniae*. In the cell mixture, rdPF4 bound predominantly to bacteria (Fig. 1E), suggesting that it has a higher affinity for bacteria than for other cell types.

**Figure 1.**
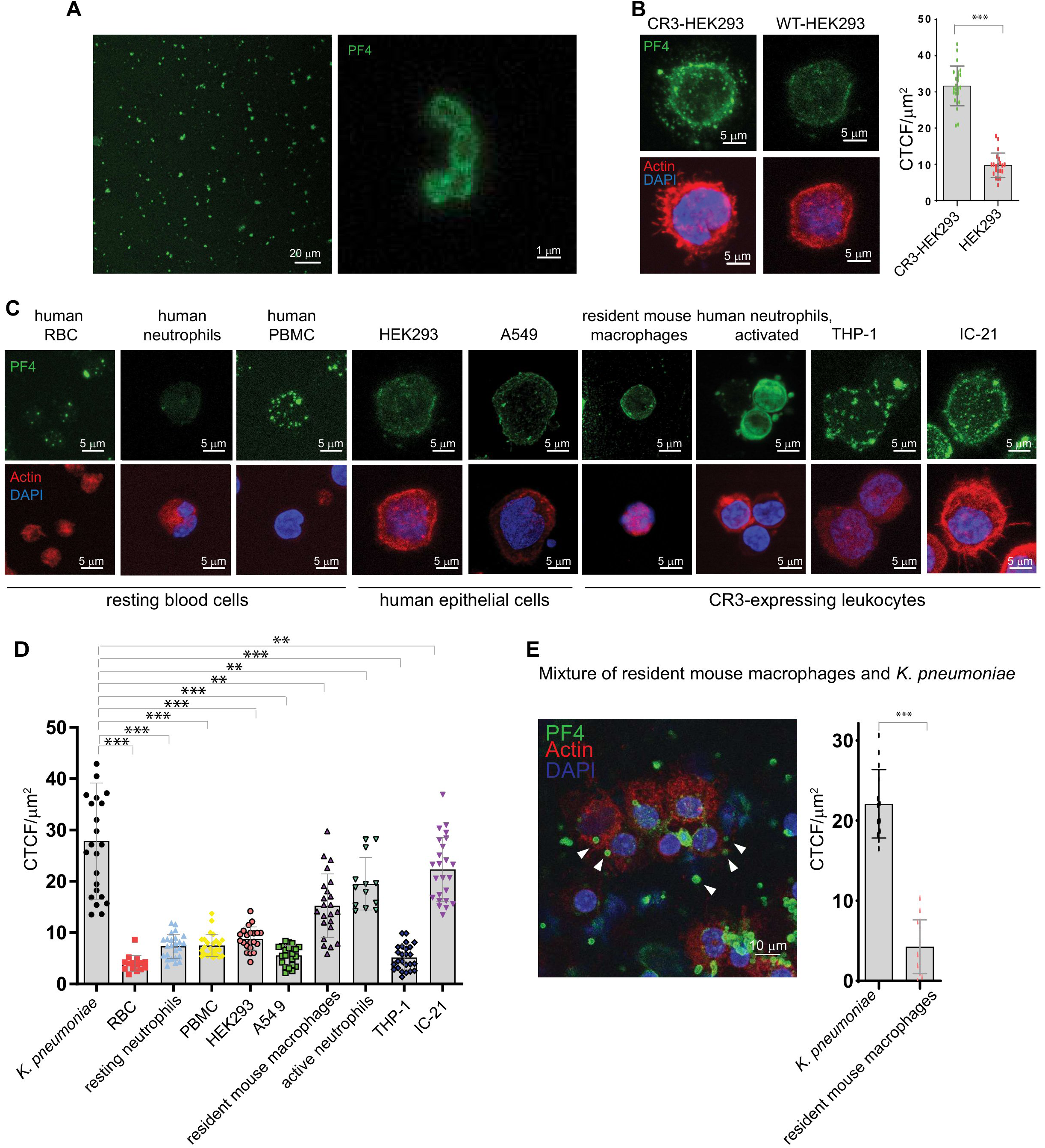
Binding of rdPF4 to *K. pneumoniae* and mammalian cells. **A**, Confocal images showing rdPF4 binding to *K. pneumoniae*. rdPF4 (100 μg/ml) was incubated with bacteria for 60 minutes at 37 °C. After removing unbound rdPF4, bacteria were incubated with rabbit polyclonal anti-PF4 antibody (1:250) for 30 minutes at 22 °C, followed by incubation with goat anti-rabbit Alexa Fluor 488-conjugated antibody. **B,** Binding of rdPF4 to the surface of CR3-expressing and wild-type HEK293 cells. Representative images are shown. Quantitation of rdPF4 surface density was performed using ImageJ. **C,** Confocal images showing rdPF4 binding (green) to various blood, epithelial, and CR3-expressing leukocytes. **D,** Quantification of rdPF4 surface density on *K. pneumoniae* and various cells from confocal images. Values are mean ± S.D. from 13 to 25 cells per cell type. **E,** Representative images of rdPF4 binding (green) to resident mouse macrophages and *K. pneumoniae* incubated in the mixture. Cells were stained with Alexa Fluor 546 phalloidin for actin and DAPI for nuclei. Arrowheads show individual bacteria. The scale bars are 20 and 1 µm (A), 5 µm (B)(C), and 10µm (E).

### Effect of rdPF4 on the growth of K. pneumoniae

Consistent with our previous findings showing that rdPF4 does not have bactericidal or bacteriostatic effects on *S. aureus* (28), incubation of *K. pneumoniae* in LB medium (pH 7.0) with increasing concentrations of rdPF4 for 3 hours did not affect bacterial growth (Fig. 2, A and B). During incubation in LB media and human plasma, bacteria-bound rdPF4 remained in mostly intact form, as shown by Western blot analyses using an anti-PF4 antibody (Fig. 2C and Fig. S1). Similarly, unbound rdPF4 in the supernatant apparently did not degrade (Fig. 2D). High-molecular-weight derivatives of bacteria-bound rdPF4 were detected after 30 min of incubation, and their amount increased after 2-3 hours, although no such products were found in the supernatant. Since bacterial growth increased several-fold during incubation with rdPF4, this indicates that new bacteria bound the free protein in solution, and that rdPF4 associated with the bacteria did not inhibit cell division.

**Figure 2.**
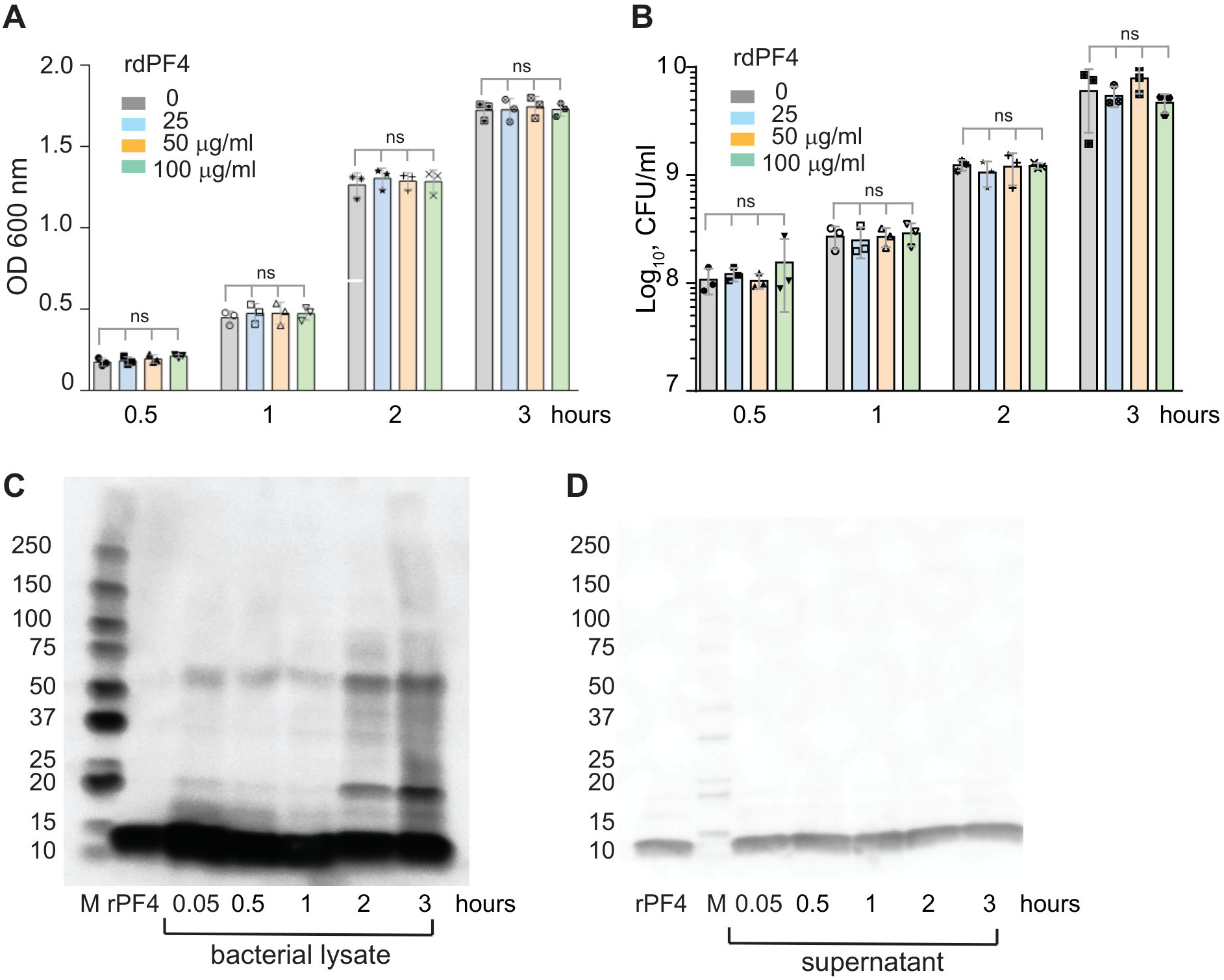
The effect of rdPF4 on bacterial growth. A, Effect of rdPF4 on the growth of *K. pneumoniae*. Bacteria were grown overnight in LB media and diluted 1:100 in fresh LB media. Bacterial suspensions (1 ml) were incubated with different concentrations of rdPF4 (25, 50, and 100 µg/ml) for 0.5-3 hours at 37 °C, and the OD at 600 nm was measured. **B,** Serial dilutions of bacteria were plated on LB agar and incubated for 24 hours at 37 °C. Colonies were counted, and results are expressed as CFU/ml. **C, D,** Stability of rdPF4 during incubation with *K. pneumoniae* was assessed by Western blotting. Bacterial suspensions were incubated with 100 µg/ml rdPF4 for 0.5-3 hours at 37 °C. Pelleted cells (**C**) and supernatants (**D**) were analyzed using polyclonal anti-PF4 antibody. M, protein standards.

### The polysaccharide capsule of K. pneumoniae prevents the binding of CR3-expressing macrophages

In its cationic ligands, including rdPF4, CR3 favors sequences enriched in positively charged residues (25). To investigate whether the antiphagocytic properties of the bacterial capsule are due to its negative charge, which could prevent contact with CR3, we tested adhesion of two representative CR3-expressing cell lines to surfaces coated with CPS isolated from the K3 serotype of *K. pneumoniae*. THP-1 macrophages and model CR3-HEK293 cells failed to adhere to plastic coated with increasing concentrations of CPS (Fig. 3A; shown for CR3-HEK293 cells). Similar to CPS, heparan sulfate, another negatively charged polysaccharide, did not support adhesion (Fig. 3A). In contrast, when immobilized directly on plastic, rdPF4 supported strong adhesion (Fig. 3A). rdPF4 also bound to immobilized CPS with an apparent *K*d of 58 nM (Fig. 3B). Treating CPS-coated surfaces with rdPF4 converted the non-adhesive CPS into an adhesive substrate able to support cell attachment (Fig. 3C). Furthermore, cells spread more effectively on rdPF4-coated CPS surfaces, indicating strong adhesion (Fig. 3, D and E). These findings suggest that CR3’s inability to bind negatively charged bacterial CPS may be a significant factor enabling *K. pneumoniae* to evade phagocytosis, and that binding of rdPF4 can disable this antiphagocytic shield.

**Figure 3.**
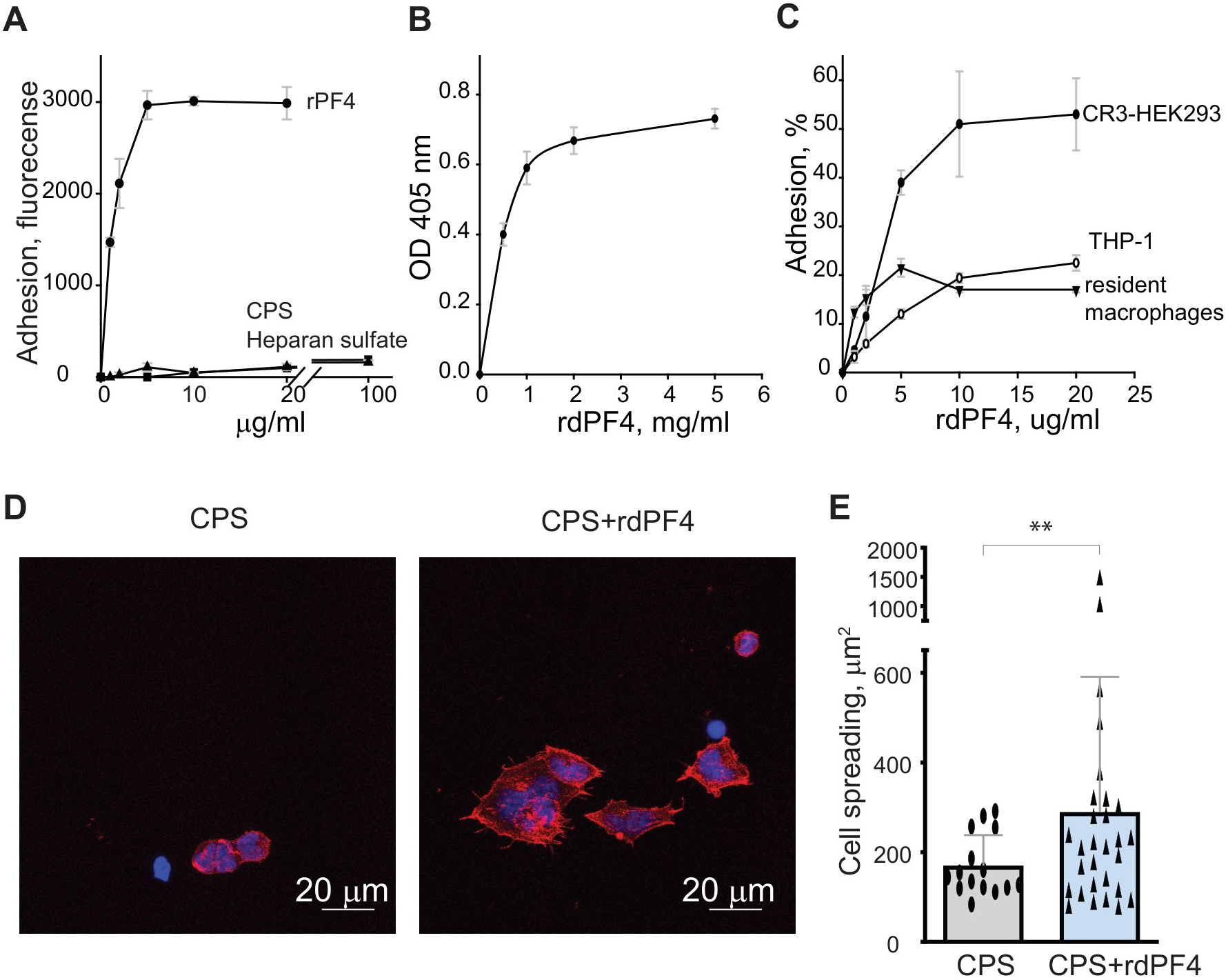
The interaction of rdPF4 with negatively charged surfaces converts them into adhesive substrates for CR3-expressing cells. **A.** Microtiter plates were coated overnight at 4 °C with varying concentrations of CPS (1-100 µg/ml), heparan sulfate (1-100 µg/ml), or rdPF4 (1-20 µg/ml). After washing with PBS, calcein-labeled CR3-HEK293 cells (5×10⁴/0.1 ml in HBSS) were added and incubated for 30 minutes at 37 °C. Nonadherent cells were removed, and the fluorescence of bound cells was measured using a fluorescence plate reader. Adhesion is expressed as the fluorescence of adherent cells, in arbitrary units. Data are normalized to adhesion on uncoated plastic (0) and are shown as means ± S.D. from three independent experiments performed in triplicate at each data point**. B,** Wells were coated with CPS, washed, and blocked with 1% PVP for 1 hour at 22 °C. Increasing concentrations of rdPF4 (0–40 µg/ml) were added for 3 hours at 37 °C, and bound protein was detected using a polyclonal anti-PF4 antibody. **C,** Wells coated with CPS and blocked with PVP were incubated with various concentrations of rdPF4 (0–40 µg/ml) for 3 hours at 37 °C. Calcein-labeled CR3-HEK293 cells, THP-1 macrophages, or mouse resident peritoneal macrophages (5×10⁴/0.1 ml) were then added. Adhesion was measured after 30 minutes at 37 °C. Results represent means ± SD from three experiments. **D,** Representative images of CR3-HEK293 cells spread on CPS-coated surfaces without rdPF4 (*left panel*) or treated with 10 µg/ml rdPF4 (*right panel*). Cells were fixed with 2% paraformaldehyde and stained with phalloidin conjugated to Alexa Fluor 568. Scale bar: 20 µm. **E,** Quantification of cell spreading. Cell area was measured in ImageJ from 25 cells per condition, across three random 20× fields. Data are presented as means ± S.D. **p < 0.01.

### rdPF4 promotes phagocytosis of heat-inactivated K. pneumoniae by CR3-expressing leukocytes

To evaluate the effect of rPF4 on phagocytosis, we initially used the K3 strain of *K. pneumoniae* (ATCC 13883), which was previously identified as a low-virulence (lv*Kp*) strain that did not cause mortality in CD1 mice infected intraperitoneally (i.p.) with 10^6^ CFU (16). The effect of rdPF4 on phagocytosis was assessed using fluorescein-labeled heat-inactivated bacteria and various CR3-expressing cultured and primary leukocytes in suspension. As measured by FACS analysis, rdPF4 dose-dependently enhanced phagocytosis of bacterial particles by a mouse peritoneum-derived macrophage IC-21 cell line (Fig. 4, A and B). At 65 μg/ml, rdPF4 increased phagocytosis by 8.6-fold. rdPF4 also significantly increased phagocytosis by different primary phagocytes (Fig. 4, C-F). However, the cell responses to rdPF4 varied. At 75 µg/ml, rdPF4 modestly (∼2.5-3.5-fold) increased phagocytosis by resident and inflammatory macrophages, as well as human neutrophils, but it boosted phagocytosis by approximately 40-fold in a mixture of neutrophils and monocytes isolated from the mouse peritoneum 4 hours after TG injection.

**Figure 4.**
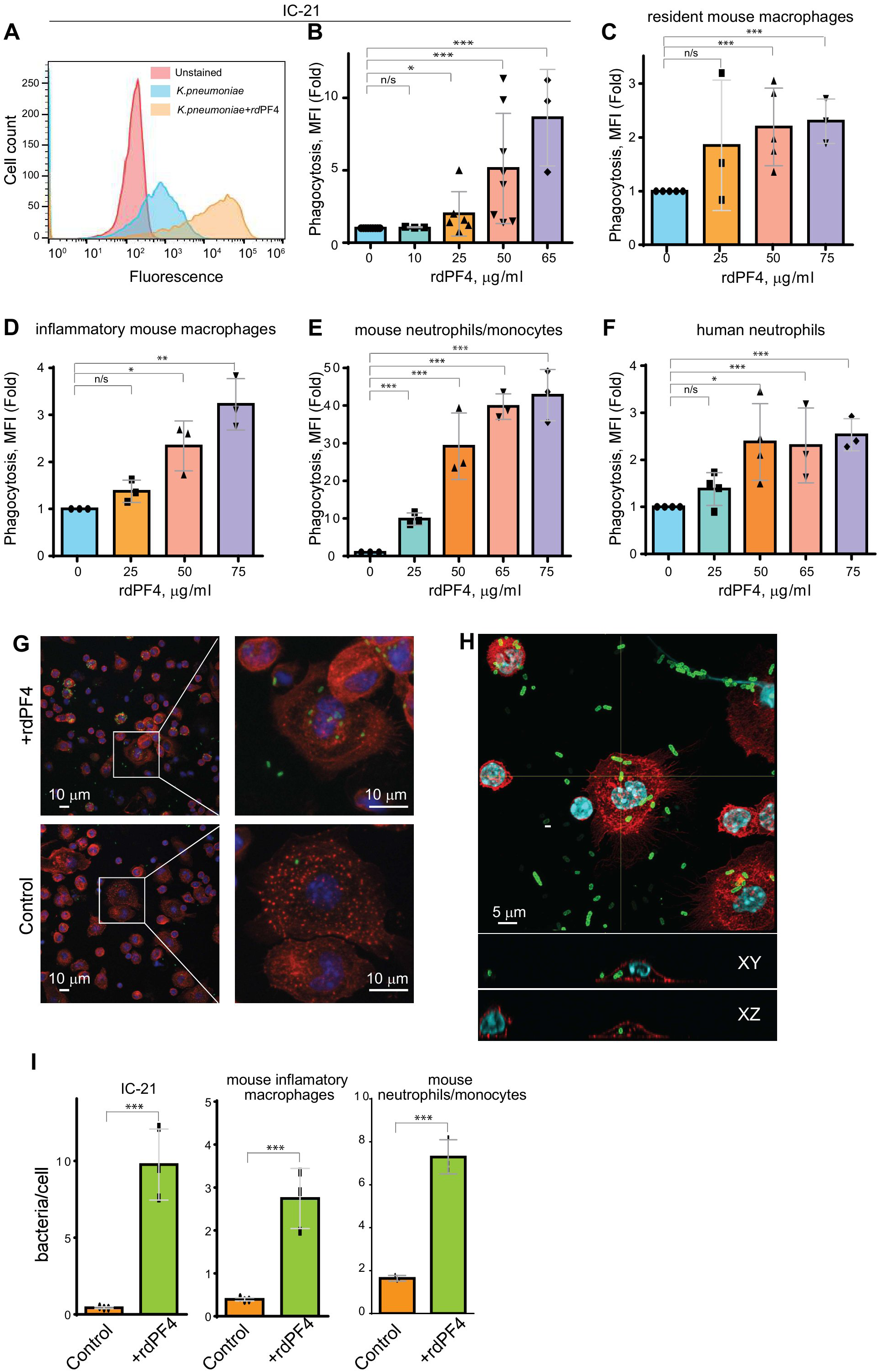
rdPF4 enhances phagocytosis of heat-inactivated *K. pneumoniae* by macrophages. **(A)** Effect of rdPF4 on phagocytosis of *K. pneumoniae* by suspended IC-21 macrophages. Cells were incubated with fluorescein-labeled, heat-inactivated *K. pneumoniae* (ATCC 13883; capsule type K3) pretreated with 50 µg/ml rdPF4 for 30 minutes at 37 °C. Phagocytosis was measured using flow cytometry. Shown are representative histograms from eight independent experiments. **(B)** Quantification of the effect of different concentrations of rdPF4 on phagocytosis of *K. pneumoniae* by suspended mouse IC-21 macrophages based on flow cytometry results. MFI, mean fluorescence intensity. Data are presented as the mean fold difference in MFI ± standard deviation from 3-8 independent experiments. *p < 0.05, *\*\**p < 0.01, *\*\*\**p < 0.001, ns, nonsignificant. **(C-F)** Quantification of phagocytosis of *K. pneumoniae* by various phagocytes in the presence of different concentrations of rdPF4. Values represent MFI fold-difference means ± S.D. from 3-8 independent experiments. **(G)** Fluorescein-labeled, heat-inactivated *K. pneumoniae* were pretreated with 50 µg/ml rdPF4 and added to adherent TG-elicited inflammatory macrophages for 60 minutes at 37 °C. Non-phagocytosed bacteria were removed, and phagocytosis was assessed from fluorescent cell images. Representative images of macrophages exposed to rdPF4-coated bacteria (*upper panel*) and uncoated control bacteria (*lower panel*) are shown. Boxes indicate enlarged regions in the right panels. The *scale bars* are 10 μm. **(H)** Representative confocal image showing phagocytosed fluorescein-labeled *K. pneumoniae* inside a macrophage. Horizontal (XY) and vertical (XZ) cross-sections of the macrophage were taken at the positions shown by white lines. The *scale bar* is 5 μm. **(I)** Quantification of rdPF4-mediated enhancement of *K. pneumoniae* phagocytosis, expressed as the number of fluorescent bacteria per cell. Phagocytosis was assessed in IC-21 macrophages, mouse inflammatory macrophages, and mouse neutrophil/monocyte populations. Values represent mean ± S.D. from three random fields per condition in three independent experiments *** p < 0.001.

Since bacterial entry into the peritoneum causes rapid macrophage adherence (37), we also examined the effect of rdPF4 on the uptake of fluorescently labeled bacterial particles by adherent macrophages. After adding rdPF4-coated bioparticles to adherent macrophages for 1 hour, their numbers inside macrophages were determined. Fig. 4G shows that rdPF4 significantly increased phagocytosis by macrophages isolated from the inflamed mouse peritoneum. To ensure that only fluorescent particles engulfed by macrophages were counted, we analyzed the stack of images for each cell, which consisted of 7-8 horizontal confocal sections (Fig. 4H). When used at 75 µg/ml, rdPF4 increased phagocytosis by 23.2 ± 7.6, 7.0 ± 2.3, and 4.3 ± 0.25-fold in IC-21 cells, mouse inflammatory macrophages, and a mixture of inflammatory neutrophils and monocytes isolated from a mouse peritoneum 4 hours after TG injection (Fig. 4I).

### rdPF4 enhances the clearance of low-virulence K. pneumoniae in the mouse sepsis model and improves survival

To determine whether the phagocytosis-promoting activity of rdPF4 observed in vitro can enhance bacterial clearance in vivo, we examined the effect of rdPF4 on lv*Kp* clearance in a mouse infection model. Previous studies showed that both i.p. and i.v. injections of *K. pneumoniae* caused pneumonia, bacteremia, and bacterial dissemination to various organs (16, 38, 39), recapitulating the features of sepsis. Therefore, we analyzed bacterial distribution in the peritoneum, blood, lungs, and liver in infected mice treated with rdPF4. Mice were infected i.p. with 1×10^8^ CFU of lv*Kp*, followed by an intraperitoneal injection of 12 µg of rdPF4 per mouse. The rdPF4 dose was selected based on our previous studies, which demonstrated that this dose significantly improved *S. aureus* clearance (28). *K. pneumoniae* counts in the peritoneal lavage and selected organs were measured 24 hours post-infection. As shown in Fig. 5, A-D, a single dose of rdPF4 significantly increased bacterial clearance by ∼22-, 10-, 2.5-, and 5.5-fold in the peritoneum, blood, lung, and liver, respectively.

**Figure 5.**
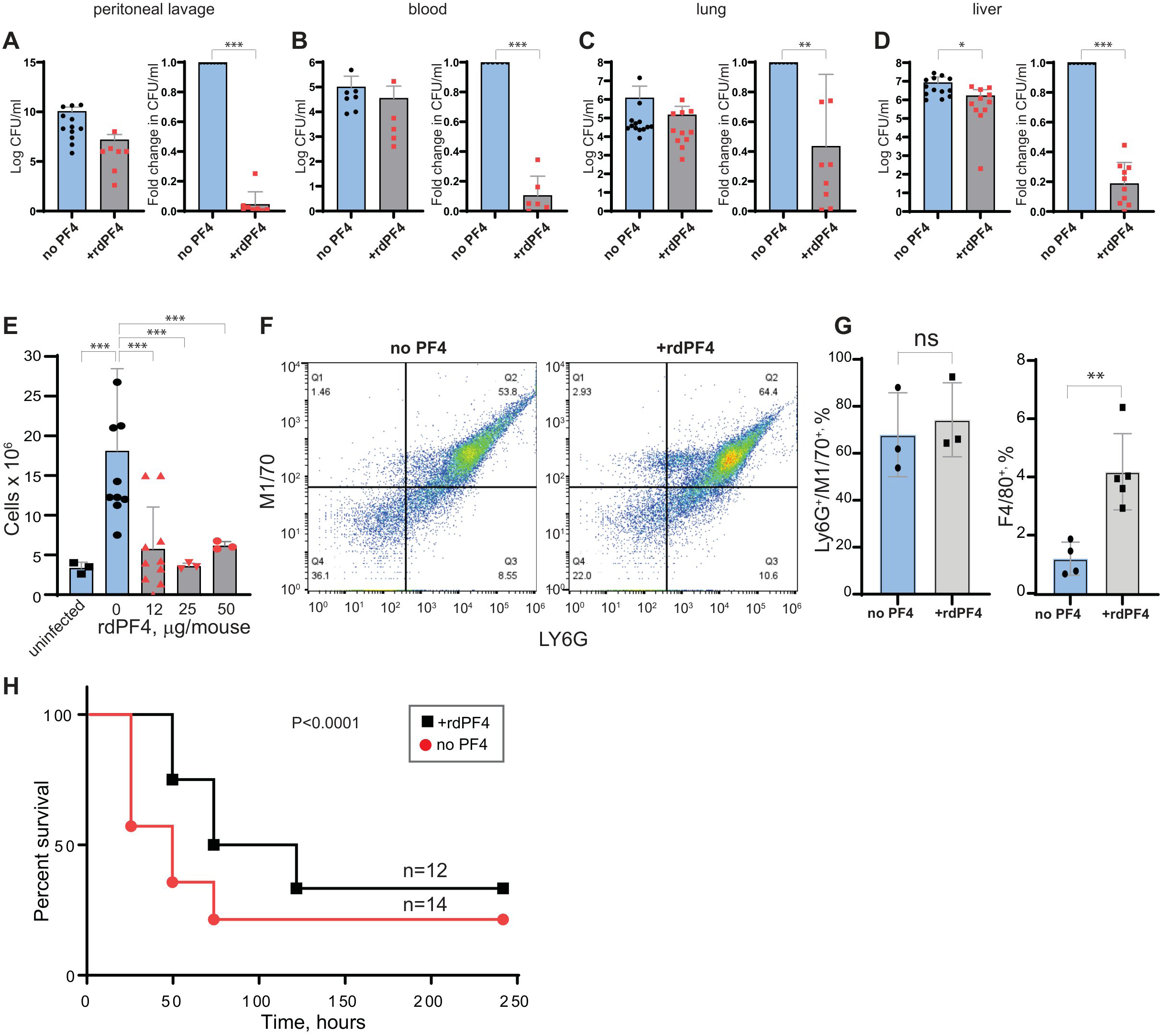
rdPF4-dependent clearance of low-virulence *K. pneumoniae* in a mouse sepsis model. (A-D) C57BL/6 mice were infected i.p. with lv*Kp* (1×10^8^ CFU) alone or with 12 µg/mouse of rdPF4. After 24 hours, samples of peritoneal lavage (A), blood (B), lung (C), and liver (D) tissue were collected. Peritoneal lavage and blood samples were serially diluted and plated on LB agar for 16 hours at 37°C. Lung and liver tissues were weighed, homogenized, and passed through a 100-μm cell strainer. The collected samples were serially diluted, plated on LB agar, and incubated for 16 hours at 37°C; colonies were then counted. Values shown are CFU/ml (*left of each panel*) and fold changes in CFU/ml means ± S.D. (*right of each panel*), from 5 to 12 mice per group. **(E)** Peritoneal lavage was collected 24 hours after injecting *K. pneumoniae* (1×10^8^ CFU) with varying concentrations of rdPF4, and the total cell count was determined. Data are presented as means ± S.D. from 3-9 mice per group. **(F)** Identification of mouse neutrophils in peritoneal lavage 24 hours after *K. pneumoniae* infection in the absence (*left panel*) or presence of rdPF4 (*right panel*). Peritoneal cells were incubated with anti-Ly6G mAb (neutrophil marker), and anti-CR3 mAb M1/70. Shown is the total population gated on single cells. (**G**) The number of neutrophils (Ly6G^+^/M1/70^+^) and macrophages (F4/80^+^) in the cell population. The data are the means ± S.D. from 3-5 experiments. **(H)** Kaplan-Meier analysis of mice (n=12-14 mice per group) after IP injection of *K. pneumoniae* (5×10^8^ CFU) without or with 25 µg/mouse (1.25 mg/kg) of rdPF4. p <0.0001.

Analysis of cell numbers in the peritoneal lavage of infected mice showed that *K. pneumoniae* caused a strong influx of leukocytes into the peritoneum (Fig. 5E). Additionally, consistent with our previous data from mice infected with *S. aureus* (28), significantly fewer leukocytes were observed in the peritoneum of infected mice after injection of 12 µg/mouse rdPF4 (Fig. 5E). Higher doses of rdPF4 did not further reduce leukocyte numbers. FACS analyses using pan-granulocyte anti-CR3 mAb M1/70, neutrophil-specific anti-Ly6G, and macrophage-specific anti-F4/80 mAbs showed that the cell composition in the lavage fluid of mice treated with rdPF4 was different than that of untreated mice (Fig. 5F). Although there was no statistically significant difference in the number of neutrophils (67 ± 17 vs 74 ± 15) in rdPF4-treated mice, the population of cells identified as F4/80^+^ macrophages increased (Fig. 5G). To assess whether rdPF4-mediated bacterial clearance improves survival, 14 mice were infected with a sublethal dose (5×10^8^ CFU) of lv*Kp*. Administering 12 µg of rdPF4 per mouse (n=12) significantly improved survival compared with untreated controls, resulting in a 12% decrease in mortality at 10 days (Fig. 5H).

Since i.p. injection of *K. pneumoniae* produces histopathological features of pneumoniae (39), we evaluated the effect of rdPF4 on histological changes in lung tissue. Consistent with the lower lung CFU count, treating mice with rdPF4 resulted in less inflammation than in untreated mice. As shown in Fig. 6A, rdPF4 provided some protection against lung tissue injury. In particular, alveolar collapse was less severe in treated mice than in untreated mice (Fig. 6, A and B; Fig. 2S). One notable feature of rdPF4-treated mice was the ∼4-fold reduction in the number of thrombi 24 hours after infection, as measured by the total area occupied by thrombi per HPF (Fig. 6C). Furthermore, the lung thrombi were smaller than those in untreated mice. The trend toward reduced thrombi was also observed at 3 and 10 days, although it did not reach statistical significance. The thickening of the alveolar septa, an important feature of tissue injury (40), was slightly but significantly less in rdPF4-treated mice at all times after infection compared to untreated mice (Fig. 6D; Fig. 3S).

**Figure 6.**
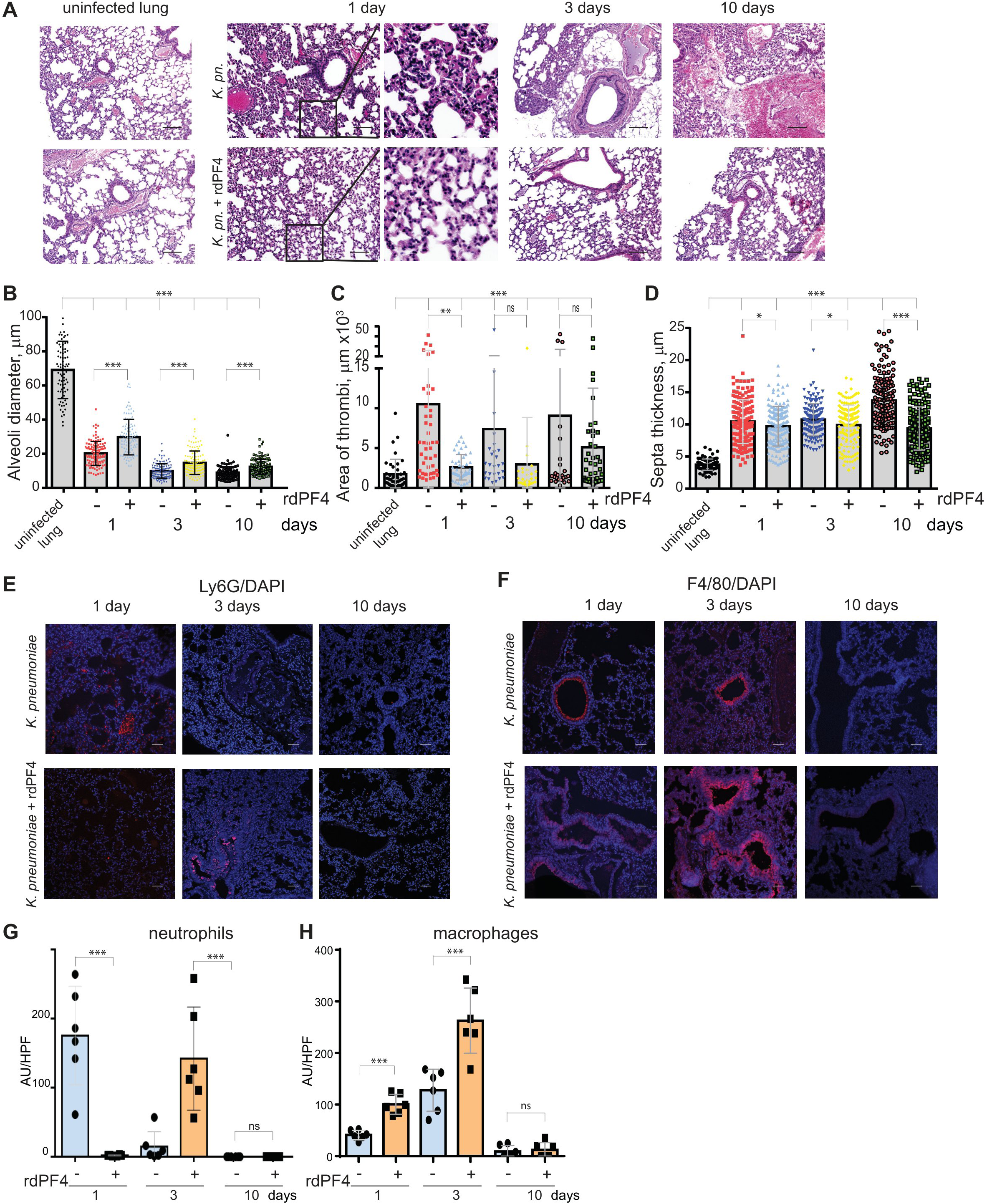
rdPF4 provides protection to lung tissue from injury caused by low-virulence *K. pneumoniae*-induced pneumonia. **(A)** Histological analyses of lung tissue isolated at 1, 3, and 10 days after infection with 1×10^8^ CFU of lv*Kp* without (upper panel) and with 12 µg/mouse rdPF4 (bottom panel). The images on the left show uninfected lung tissues. The scale bar is 100 μm. **(B)** Alveoli diameter, **(C)** thrombi area, and **(D)** septa thickness were measured from histological images. Results are presented as mean ± SD from five randomly selected images from three different samples. ***p < 0.001; ns nonsignificant. **(E, F)** Immunofluorescence images of lung tissue collected from infected (upper panel) and rdPF4-treated mice (bottom panel) 1, 3, and 10 days after infection. Tissue sections were labeled with anti-Ly6G (neutrophil marker) and DAPI **(E**) and anti-F4/80 (macrophage marker) and DAPI **(F)**. Representative images are shown. The scale bar is 50 μm. **(G, H)** Relative expression levels of Ly6G on neutrophils and F4/80 on macrophages were determined from six random fields. Data are mean ± SD. \*\*\**p* < 0.001; ns, nonsignificant.

Infection with *K. pneumoniae* is known to induce infiltration of neutrophils and monocyte/macrophages into the alveolar space and interstitial septa (1, 3). To evaluate leukocyte infiltration following rdPF4 treatment, lung tissue sections were stained with the neutrophil-specific mAb Ly6G and the macrophage-specific mAb F4/80. The tissues of untreated mice showed significant neutrophil infiltration 24 hours after infection, with neutrophils primarily present in the interstitium. The number of neutrophils decreased sharply by day 3 and stayed low on day 10 (Fig. 6, E and G). Neutrophil influx into the lungs of rdPF4-treated mice was suppressed after 24 hours, increased on day 3, and then declined. The course of F4/80-positive cell infiltration was different. In both treated and untreated mice, the number of monocytes/macrophages increased gradually from day 1 to day 3, then decreased by day 10 (Fig. 6, F and H). However, the flux of these cells in the rdPF4-treated mice was 2.5 ± 1.1 and 2.0 ± 0.52 times higher than in untreated mice on days 1 and 3. Since inflammatory monocytes/macrophages have been implicated in protection against *K. pneumoniae* (41), (16, 42), their higher proportion in rdPF4-treated mice may have contributed to the more rapid bacterial clearance.

To assess how neutrophils and monocytes/macrophages contribute to the clearance of lv*Kp*, we depleted neutrophils using mAb 1A8, which specifically targets neutrophils, and measured bacterial clearance in the peritoneum, lung, liver, and blood of both untreated and rdPF4-treated mice. The effectiveness of neutrophil depletion was confirmed by the nearly complete absence of Ly6G-positive cells recruited to the peritoneum 6 hours after TG injection (Fig. 7, A and B). Neutrophil depletion in mice infected with 3×10^7^ CFU lv*Kp* led to higher bacterial counts in the depleted mice compared to nondepleted mice (75 ± 21, 109 ± 64, 27 ± 3, and 2135 ± 2266-fold in the peritoneal lavage, lungs, liver, and blood, respectively) 24 hours post-infection, highlighting the importance of neutrophils in bacterial clearance (Supplemental Table). The lack of neutrophils particularly impaired bacterial clearance in the blood and, to a lesser degree, affected the liver, peritoneum, and lungs. Treatment with rdPF4 (12 µg/mouse) reduced bacterial loads in organs and fluids by approximately 3-4-fold (Fig 7C). Furthermore, a single treatment with 12 µg of rdPF4 per mouse (n=8) significantly improved survival compared to untreated controls (n=6), suggesting that remaining monocytes/macrophages might have contributed to bacterial clearance (Fig.7D). Depletion of monocytes and macrophages using Clodronate liposomes caused a significant reduction in macrophages in mice infected with 1×10^7^ CFU lv*Kp* (as confirmed by the absence of F4/80-positive cells in the peritoneum) and resulted in higher bacterial counts compared to nondepleted mice (Supplemental Table). Except for the lungs, macrophage depletion strongly impaired bacterial clearance in the peritoneum (∼10^4^-fold) and the liver (∼2 × 10^3^-fold) compared to nondepleted mice, and this impairment was much greater than in neutrophil-depleted mice, even though a lower dose of lv*Kp* was used for infection. Monocyte depletion affected bacterial clearance in the blood to a similar extent as neutrophil depletion. Surprisingly, bacterial clearance in the lungs was significantly less affected than in other organs and to a lesser extent than in neutrophil-depleted mice. Furthermore, there was no significant difference in bacterial counts across all organs and fluids of rdPF4-treated and untreated macrophage-depleted mice, indicating that macrophages are essential for the effect of rdPF4.

**Figure 7.**
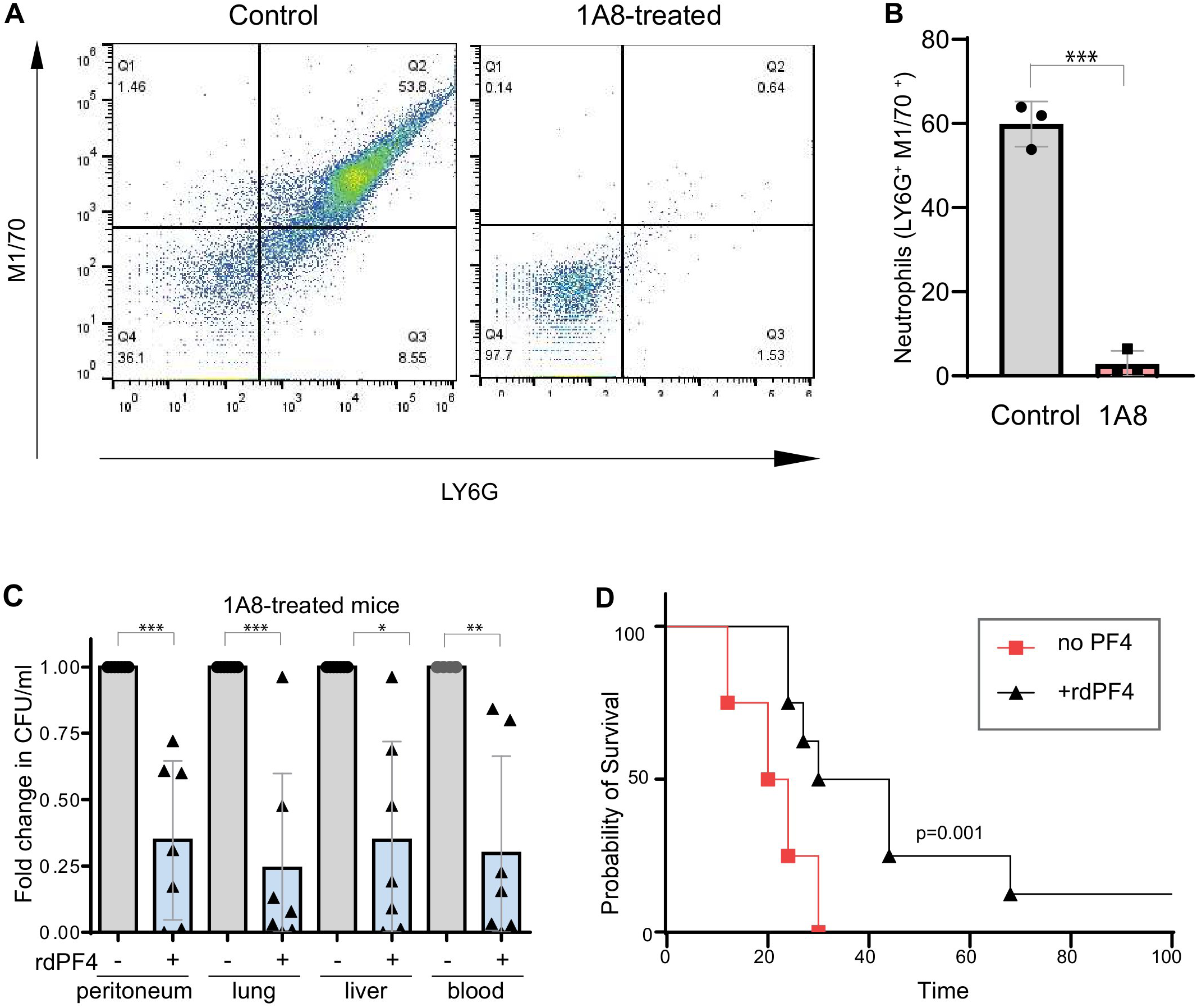
Effect of rdPF4 on clearance of low-virulence *K. pneumoniae* in neutrophil-depleted mice. **(A, B)** To confirm neutrophil depletion, C57BL/6 mice were injected with anti-Ly6G mAb 1A8 (100 µg/mouse) and, after 24 hours, injected with TG to induce the neutrophil flux into the peritoneum. Peritoneal lavage was isolated 6 hours after TG injection, and cells were analyzed by FACS using anti-Ly6G mAb and anti-CR3 mAb M1/70; ***p < 0.001. **(C)** Mice (n=6 per group) were treated for 24 hours with 100 µg of mAb 1A8, and then injected with 3×10^7^ lv*Kp* and rdPF4 (12 µg/mouse). The effect of rdPF4 on bacterial clearance in the peritoneum, blood, lungs, and liver in neutropenic mice was determined 24 hours after infection. *p < 0.05, *\*\**p < 0.01, *\*\*\**p < 0.001. **(D)** Kaplan-Meier analysis of neutrophil-depleted mice (n=6-8 mice per group) after i.p. injection of lv*Kp* (5×10^7^ CFU) without or with 12 µg/mouse (0.6 mg/kg) of rdPF4. p =0.001. For survival experiments, mice were administered 100 µg of mAb 1A8 24 hours before lv*Kp* infection, followed by repeated doses of 100 µg every 48 hours thereafter.

### Effect of rdPF4 on phagocytosis of high-virulence K. pneumoniae and its clearance during septic infection

The commonly studied ATCC 43816 K2 strain of *K. pneumoniae* was previously identified as high-virulence (hv*Kp*), with a mean survival time of 12 hours in CD1 mice infected i.p. with 10^6^ CFU (16). Similar to its low-virulence counterpart, rdPF4-coated hv*Kp* was more susceptible to phagocytosis than untreated bacteria. As shown in Fig. 8 (A-D and I), rdPF4 enhanced phagocytosis of pHrodo-labeled heat-inactivated bacteria by various phagocytes in a dose-dependent manner. rdPF4 also significantly increased the phagocytosis of live bacteria; however, its effect was smaller than with dead bacteria (Fig. 8, E-H and I).

**Figure 8.**
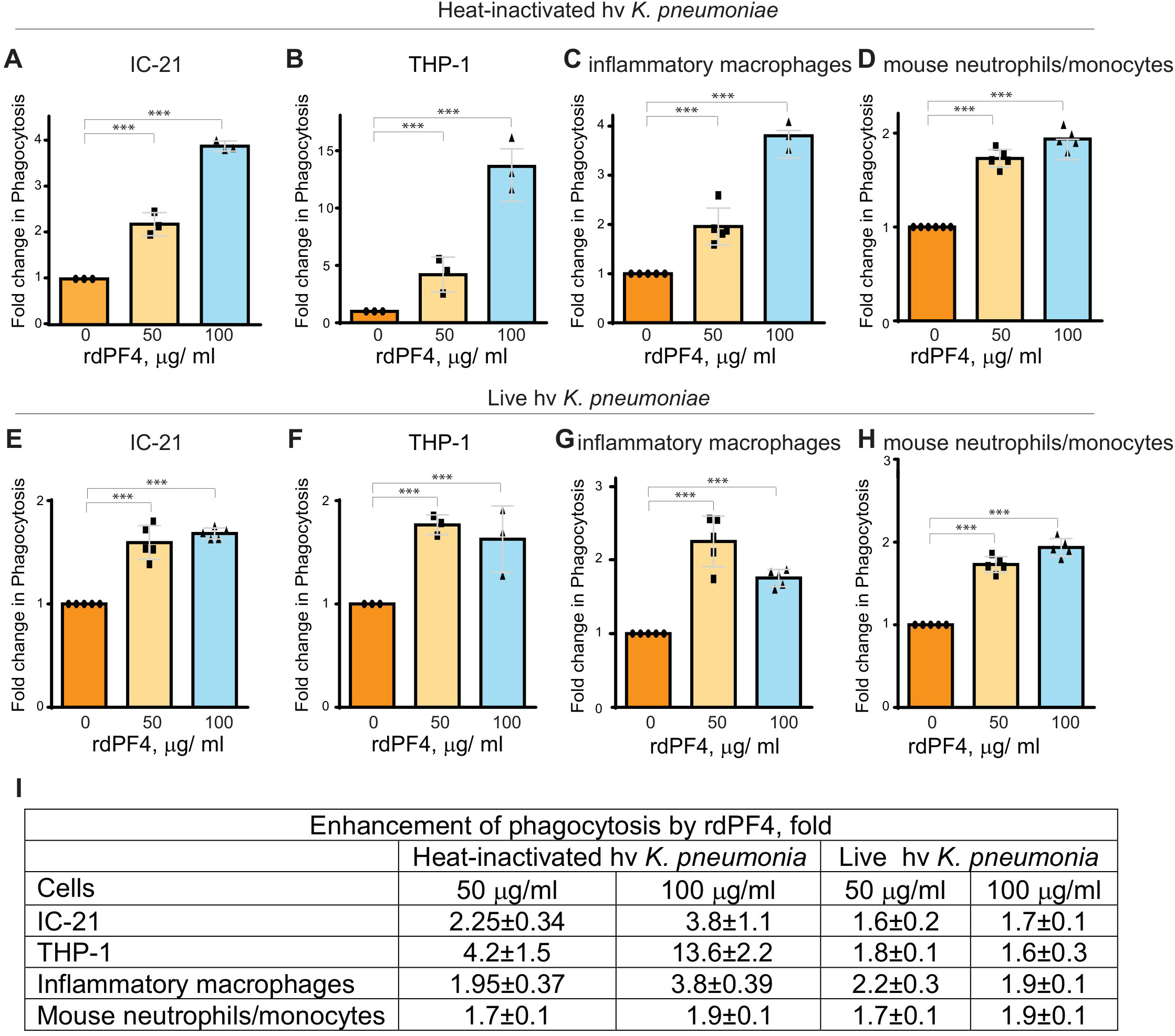
rdPF4 enhances phagocytosis of heat-inactivated and live high-virulence *K. pneumoniae* by macrophages. **(A-D)** Heat-inactivated pHrodo-labeled hv*Kp* (ATCC 43816; capsule type K2) were preincubated with various concentrations of rdPF4 for 1 hour at 37 °C, and rdPF4-coated bacteria were then added to suspended IC-21 macrophages, PMA-activated THP-1 cells, TG-elicited inflammatory mouse macrophages, and mouse peritoneal neutrophils/monocytes for 30 minutes at 37 °C, and phagocytosis was measured by flow cytometry. **(E-H)** Live bacteria were labeled with pHrodo, preincubated with rdPF4, and then added to various leukocyte populations to assess phagocytosis. Data are presented as means ± S.D. from three separate experiments. ***p< 0.001. **(I)** The table shows fold changes in phagocytosis at two concentrations of rdPF4 (50 and 100 μg/ml) relative to phagocytosis in the absence rdPF4 across various macrophage types.

The effect of rdPF4 on bacterial clearance was examined in mice infected i.p. with 10^4^ CFU of hv*Kp*. A lower bacterial dose was chosen to prolong the survival time of C57BL/6 mice to approximately 24 hours. As shown in Fig. 9A, a single dose (12 µg/mouse) of rdPF4 improved bacterial clearance, reducing the number of live bacteria in the peritoneal lavage, blood, lung, and liver by 12.3 ± 9.1, 6.6 ± 5.0, 5.0 ± 3.8, and 4.3 ± 3.4-fold, respectively.

**Figure 9.**
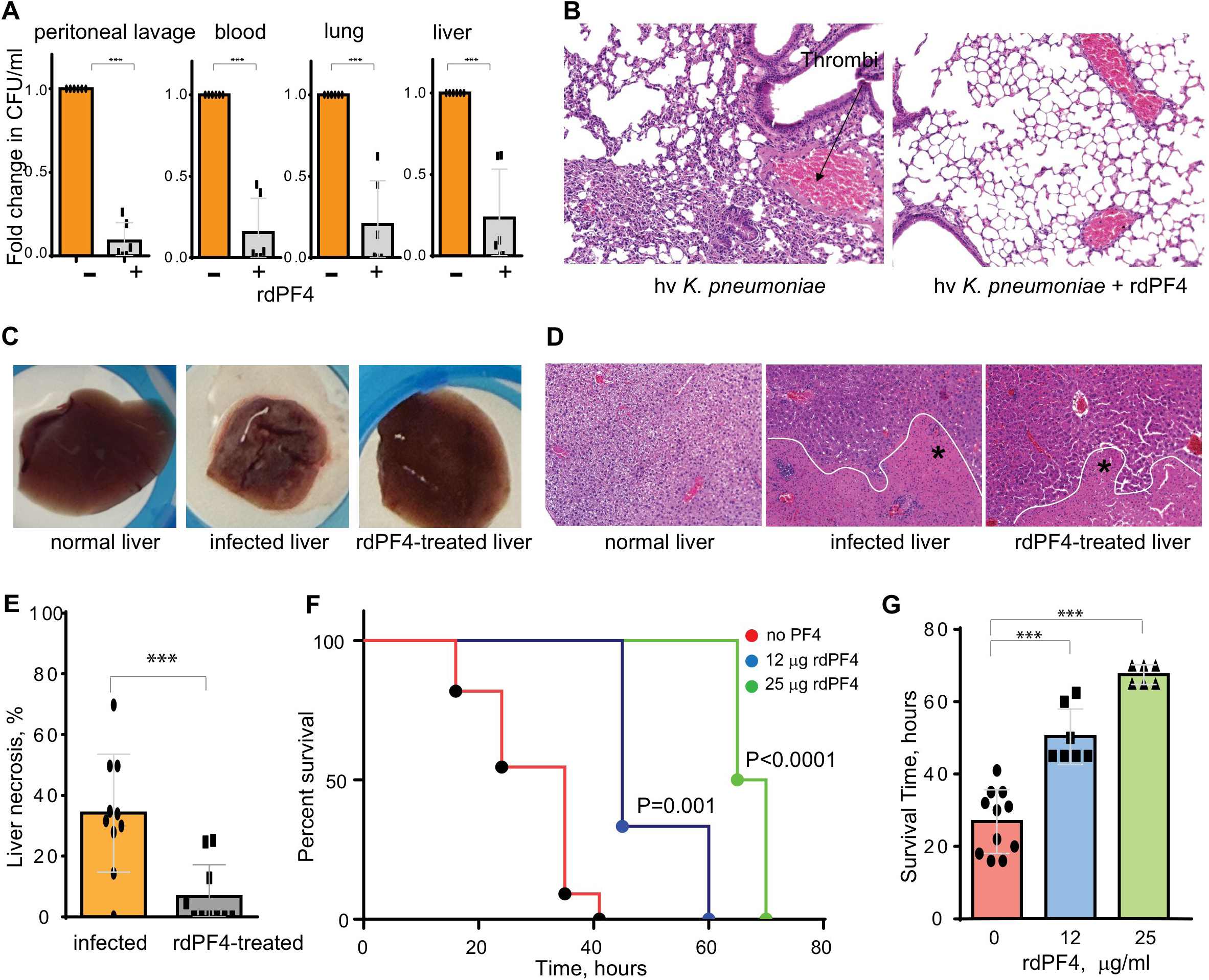
rdPF4 enhances the clearance of high-virulence *K. pneumoniae*, reduces tissue injury, and improves mouse survival rates. **(A)** Bacterial loads in peritoneal lavage, blood, lung, and liver were collected 24 hours after infection of C57BL/6 mice with an hv*Kp* (10⁴ CFU). Data show the fold change in bacterial load (CFU/ml) between rdPF4-treated and untreated mice (6 mice per group), expressed as mean ± S.D. **(B)** H&E-stained sections were collected from the lungs of control (*K. pneumonia* only) and rdPF4-treated mice 24 hours after infection. **(C)** Representative images of liver lobes from uninfected, infected, and rdPF4-treated mice 24 hours after infection. **(D)** H&E-stained liver sections from normal, infected, and rdPf4-treated mice. **(E)** Quantification of necrotic areas in liver sections. Values represent mean ± S.D. from 10 images per group. **(F)** Kaplan–Meier survival curves of mice (n = 6-10 per group) following intraperitoneal infection with hv*Kp* (3×10⁴ CFU) alone or treated with rdPF4 at different concentrations. **(G)** Survival time of mice infected with hv*Kp* and treated with varying doses of rdPF4. Data represent mean ± S.D. from groups of 6-10 animals. ***p < 0.001.

Histological analyses revealed that, similar to mice infected with a high dose of lv*Kp*, the hvKp strain caused severe lung inflammation 24 hours after infection, with many areas displaying collapsed alveoli, thickened septa, and thrombi (Fig. 9B). Additionally, bronchioles in many areas showed edema, and blood vessels exhibited endothelial cell degeneration and loss of the basement membrane. When treated with rdPF4, the lungs of animals challenged with hv*Kp* demonstrated more normal morphology (Fig. 9B).

Capsular serotypes K1 and K2 of *K. pneumoniae* have been identified as the main virulence factors associated with severe complications, especially liver abscesses (4, 20, 43, 44). As shown in Fig. 9C, compared to uninfected animals, mice infected with hv*Kp* (K2 serotype) developed multiple hepatic microabscesses as early as 24 hours after infection, and treatment with rdPF4 restored the liver tissue to a nearly normal appearance. Histological examination of the livers from hv*Kp*-infected mice revealed significant pathological changes, including dilated hepatic sinusoids and neutrophil infiltration in the perivascular regions, compared with normal tissue (Fig. 9D). Additionally, necrosis of the liver parenchyma was observed in multiple areas. Mice treated with rdPF4 showed less severe liver damage, with many areas maintaining normal sinusoid structure and decreased liver necrosis (Fig. 9, D and E).

The improved bacterial clearance and reduced organ injury apparently led to a dose-dependent increase in survival rates in mice treated with rdPF4 (Fig. 9, F and G). All mice infected with hv*Kp* alone (n=11) died within approximately 40 hours after infection, with a mean survival time of 27 ± 9 hours. rdPF4 at a dose of 12 µg/mouse (n=7) extended survival to 48 ± 6 hours, and 25 µg/mouse (n=6) increased it further to 67.5 ± 3 hours. Raising rdPF4 to 50 µg/ml did not provide additional survival benefit, although one of the six mice survived until the end of the 10-day study.

### rdPF4 is effective against antibiotic-resistant K. pneumoniae

To assess whether treatment with rdPF4 enhances the clearance of antibiotic-resistant bacteria, mice were infected with the carbapenem-resistant *K. pneumoniae* ATCC strain BAA-1705. This strain was resistant to many of the antibiotics tested, although it was partially susceptible to meropenem and tetracycline at concentrations above 2 µg/ml (Fig. 4S). Twenty-four hours after infecting the mice with 2×10^7^ CFU followed by rdPF4 injection (50 µg/mouse), tissue homogenates, blood, and peritoneal lavage were collected and plated on agar for CFU counts. rdPF4 enhanced bacterial clearance in the peritoneum, blood, lung, and liver by 5.8 ± 5, 29.9 ± 42, 15.6 ± 24, and 17.8 ± 23-fold, respectively (Fig. 10A). At the same time, at 1 µg/ml, meropenem was ineffective in clearing bacteria from the peritoneum and blood, although it showed a tendency to reduce bacteria in the lung and liver by ∼20-25%, but this trend did not reach statistical significance. Consistent with the effect of rdPF4 on bacterial clearance, the survival rates of rdPF4-treated mice infected with 5×10^7^ CFU were higher compared to untreated animals (Fig. 10, B and C). No synergistic effect was observed from combining rdPF4 and the antibiotic.

**Figure 10.**
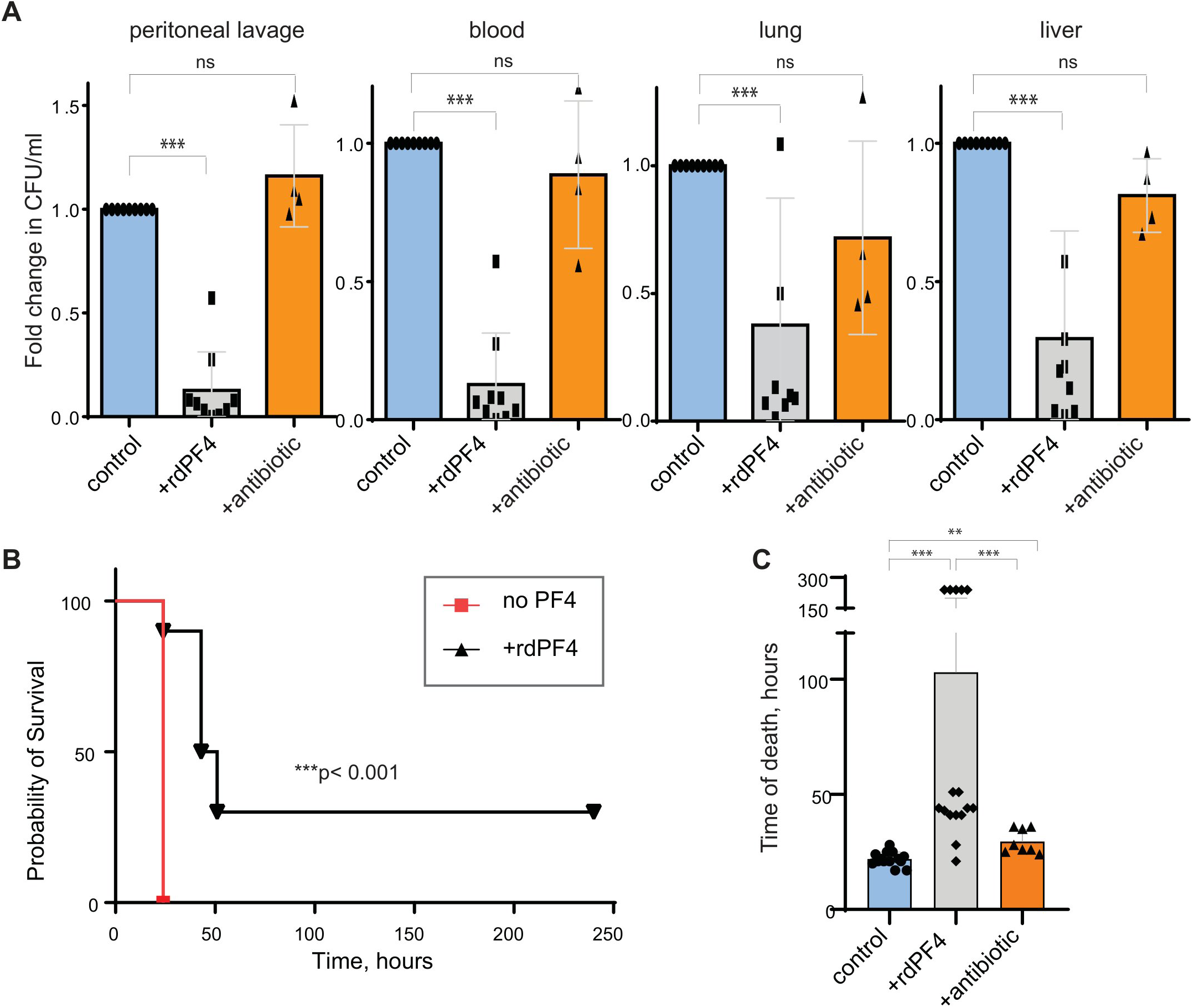
rdPF4 enhances bacterial clearance and improves survival rates in mice infected with carbapenem-resistant *K. pneumoniae* strain BAA-1705. **(A)** Mice were infected IP with the BAA-1705 strain (2×10^7^ CFU) and treated with either 50 μg/mouse (2.5 mg/kg) of rdPF4 or 125 mg/kg of meropenem. Twenty-four hours after infection, samples of peritoneal lavage, blood, lung, and liver were collected. Tissue homogenates and lavage samples were serially diluted and plated on agar for CFU enumeration. Values represent the fold differences in CFU/ml between control and treated mice (means a± S.D., n = 4-8 animals. **(B)** Kaplan-Meier analysis of mice after IP injection of BAA-1705 strain (5×10^7^ CFU) without (n=6) or with 50 μg/mouse (2.5 mg/kg) of rdPF4 (n=10). **(C)** Comparison of survival time between mice treated with rdPF4 and meropenem infected with BAA-1705. Values are mean ± S.D. from 8-16 animals per group. **p < 0.01; ***p < 0.001

## Discussion

In this study, we expand on the idea that targeting the bacterial capsule with a positively charged CR3-binding protein is an effective means of overcoming its antiphagocytic properties. We previously showed that a small cationic protein, rdPF4, a ligand for the phagocytic receptor CR3, increased phagocytosis of both nonencapsulated and encapsulated Gram-positive *S. aureus* and was effective against an MRSA strain (28). This activity led to greater bacterial clearance and improved survival in a mouse model of peritonitis. Both in vitro and in vivo experiments reported in this study demonstrate that binding of rdPF4 to low-virulence and high-virulence as well as carbapenem-resistant *K. pneumoniae* strains enhances phagocytosis by neutrophils and macrophages and improves bacterial clearance in a mouse sepsis model. These results indicate that rdPF4 has broad-spectrum activity against both Gram-positive and Gram-negative bacteria and is effective against antibiotic-resistant pathogens.

The capsule of both Gram-positive and Gram-negative bacteria has long been recognized as a key virulence factor. Encapsulated *K. pneumonia* are responsible for some of the most severe infections, including septicemia, pneumoniae, and meningitis (45–47). During bacterial infections, a dense layer of CPS forms an effective physical barrier that resists complement-mediated killing by preventing MAC assembly and blocking its ability to breach the bacterial membrane (1, 3). Numerous studies have demonstrated that while serum-resistant strains of *K. pneumoniae* can deposit the C5b-C9 complement components, they fail to mediate MAC-induced lysis (48–50). For instance, Merino et al. proposed that complement deposition occurs too far from the bacterial membrane, which prevents the formation of the lytic C5b-C9 MAC (48). Additionally, the CPS helps bacteria evade the immune system is by counteracting complement-mediated phagocytosis through masking C3b and iC3b opsonins, preventing their binding to CR3 on phagocytes (51, 52). On this point, the deposition of C3b has been shown to depend on the thickness of *K. pneumoniae* CPS (53, 54).

Along with complement resistance, the CPS, composed of negatively charged polysaccharides with many carboxyl and phosphate groups (6), has been suggested to repel phagocytes that are coated with a negatively charged glycocalyxs, thereby enhancing the CPS’s anti-phagocytic properties (55). Supporting this intuitive idea, Li et al. demonstrated that more electronegative capsules were more resistant to phagocytosis (56). Although widely accepted, the exact mechanism underlying this protective effect remains poorly understood. For instance, this model fails to explain the observation that simply neutralizing the bacterial negative charge with poly-L-lysine does not improve phagocytosis (57). In contrast, positively charged molecules containing CR3-binding sites can significantly increase phagocytosis. We previously demonstrated that rdPF4 and LL-37 bind to *S. aureus* and *E. coli* and serve as effective opsonins (30, 57). Our present data demonstrate that positively charged rdPF4 (+4.6) also binds to the CPS of *K. pneumoniae* at high density, creating multiple CR3-binding sites. Notably, rdPF4 has a relatively low molecular weight (7.8 kDa for its monomeric form and about 16 kDa for the dimer) compared to the high-molecular-weight C3b/iC3b proteins (∼185 kDa), which should facilitate deposition of many molecules of rdPF4 on the surfaces of CPS. Furthermore, CR3-binding molecules like C3b/iC3b and the iC3b-derived C3dg carry a negative charge (58) and are therefore less likely to be attracted to the negatively charged CPS. It has been documented that hv*Kp* strains, including K2 used in this study, produce a hypercapsule characterized by a thick polysaccharide layer that is more robust than a typical capsule and exhibits the hypermucoviscous phenotype (1). The mucoviscous capsule in *K. pneumoniae* is enriched in polysialic acids, which have been linked to resistance to neutrophil phagocytosis (59). Along with highly acidic glycosaminoglycans, increased polysialylation should increase the charge density of the mucoviscous CPS. Furthermore, although the CPS and mammalian glycocalyx share some common molecules, including sialic acid, bacterial capsules are generally more negatively charged and have a higher charge density than mammalian cells, which could make them preferred targets for rdPF4. Indeed, we found that the density of rdPF4 deposited on the surface of *K. pneumoniae* was approximately 3-6 times higher than that on resting primary and cultured blood cells and cultured epithelial cells (Fig. 1). Therefore, due to its cationic nature and ability to bind CPS at high density, rdPF4 not only neutralizes CPS’s negative charge but also provides numerous CR3-binding sites, thereby promoting phagocyte attraction. Additionally, unlike C3bi, which targets the outer membrane beneath the CPS, it is advantageous for rdPF4 to accumulate in the CPS’s superficial layer instead of sinking into it.

Our previous findings with *S. aureus* (28) and the results presented in this study demonstrate that rdPF4 functions as an opsonin for both Gram-positive and Gram-negative bacteria. In Gram-positive bacteria, opsonophagocytosis rather than lytic killing is the main process mediated by the complement system. Although Gram-positive bacteria can assemble the MAC on their surface, the thick peptidoglycan layer, which acts as a physical barrier, prevents MAC components from reaching the cytoplasmic membrane and causing lytic death (60). Conversely, the MAC can kill Gram-negative bacteria by forming pores in the outer membrane, since the thin peptidoglycan layer does not prevent MAC protein deposition. Nonetheless, in both bacterial groups, the CPS limits the impact of the complement system. It is well documented that not only the CPS of *K. pneumoniae* but also the capsules of *S. aureus* and *S. streptococcus* deposit C3b/iC3b underneath the capsule, shielding it from recognition by phagocytes (51). Consequently, rdPF4-mediated phagocytosis may act as an additional host defense mechanism that circumvents the complement resistance of encapsulated Gram-positive and Gram-negative bacteria, which could be particularly beneficial in mixed polymicrobial infections.

Our experiments with lv*Kp* support previous research showing that infection with *K. pneumoniae* elicits a strong inflammatory response, characterized by infiltration of neutrophils and monocytes/macrophages into the peritoneum and lungs, with neutrophils being the predominant cell type obserbed after 24 hours (Figs. 5G and 6G). Treatment with rdPF4 altered the composition of inflammatory cells, albeit in different ways across locations. While neutrophil numbers did not change in the peritoneum, the number of monocytes/macrophages was approximately 4-fold higher in rdPF4-treated mice (Fig. 5G). In the lungs, rdPF4 significantly attenuated neutrophil recruitment after 24 hours, with monocytes/macrophages being a dominant population at that time (Fig. 6H). By day 3, the recruitment of monocytes and macrophages in the lungs continued to increase, and neutrophils were also present. Although the lv*Kp*-infected liver was not examined because no significant morphological changes were observed, data from the peritoneum and lungs suggest that rdPF4 primarily increases monocyte/macrophage infiltration. Whether CCR2+ inflammatory monocytes, which have been shown to be essential for bacterial clearance in the lungs of mice infected with *K. pneumoniae* ST258 strain by intratracheal instillation (42), contributed to the increased level of F4/80-positive cells in rdPF4-treated animals remains to be determined. Also, the M1/M2 phenotype of macrophages, which has been reported to affect bacterial clearance in *Klebsiella*-infected mouse models (61–63) requires further investigations. Nonetheless, it is appealing to speculate that the less severe lung damage observed in rdPF4-treated animals infected with lv*Kp* and hv*Kp*, evident as early as 24 hours after infection and persisting through day 3 and longer (Fig. 6A and 9B), can be attributed to increased monocyte/macrophage recruitment. The lack of neutrophils observed at that time in lv*Kp*-infected lungs further supports the role of monocytes and macrophages in tissue protection. Furthermore, the finding that rdPF4 remained effective when injected into neutrophil-depleted mice suggests that monocytes and macrophages are its primary targets. However, consistent with previous studies (2, 3), neutrophils contribute to bacterial clearance, as neutrophil depletion led to less effective clearance, although to a lesser extent than depletion of monocytes/macrophages (Supplemental Table). Most notably, our finding that rdPF4 significantly protected the liver from abscesses caused by hv*Kp* can tentatively be linked to Kupffer cells, which are the largest group of tissue-resident macrophages in the body.

Our recombinant dimeric PF4 mimics half of the tetrameric protein stored in mammalian platelet α-granules at unusually high concentrations. PF4 is released from activated platelets at sites of vascular injury and is primarily trapped within platelet-rich blood clots, where its concentration was estimated at 280 μM, more than 100 times higher than serum levels (64). Although PF4 has been assigned to the CXC chemokine subfamily, it lacks a typical N-terminal ELR motif necessary for binding to G-protein-coupled chemokine receptors. In vitro studies showed that although PF4 can induce leukocyte migration, these effects require micromolar concentrations of rdPF4 (65), (64), (30), which are uncommon for chemokine-mediated GPCR activation. We recently identified CR3 (Mac-1, integrin αMβ2) as the primary receptor for rdPF4 and showed that the CR3-rPF4 interaction supports a potent migratory response in neutrophils and macrophages, entirely dependent on CR3 (30). CR3 is a multiligand receptor and its characteristic feature is the ability to bind not only the complement fragment iC3b (66, 67) but also many cationic proteins and peptides through sequences enriched in positively charged and hydrophobic amino acid residues (26, 29). CR3 recognition sequences do not have a specific consensus motif; instead, they feature motifs with cores of positively charged residues, which are flanked by hydrophobic residues (25). The identification of PF4 as a CR3 ligand was based on its cationic nature and the presence of several sequences, such as ^12^CVKTTSQ<u>VRPRHI</u>TS^26^ and ^57^<u>APLYKKIIKKLL</u>ES^70^ favored by CR3 (30). The CR3 recognition sequences are found in many cationic proteins and peptides that are stored in neutrophil and platelet granules or, like cathelicidin LL-37 and defensins, are also produced in epithelial cells and released at sites of infection or injury (57). Additionally, several examined cationic peptides from different species contain CR3 recognition motifs, including bovine bactenecin 5, Drosophila drosocin, porcine cathelicidin-derived tritrpticin, and others, and can interact with the ligand-binding αMI-domain of CR3 (25). All of these molecules belong to a broad group of antimicrobial peptides (AMP) that kill bacteria and other pathogens by disrupting or penetrating microbial membranes, which leads to cell lysis (68). Since most naturally occurring AMPs have a net positive charge (usually between +2 and +11) and a significant portion of hydrophobic regions, they can serve as potential CR3 ligands. Furthermore, although bacteriostatic, microbicidal, and lytic properties of AMPs are well documented, they can also enhance CR3-mediated phagocytosis. In this regard, we and others have demonstrated that the human cathelicidin peptide LL-37, one of the best-studied peptides in this group, is a CR3 ligand and a potent bacterial opsonin (57, 69). Tachyplesin III, an antimicrobial peptide from horse show crab, has also been shown to promote macrophage phagocytosis (70). Whether the antimicrobial opsonic activity of other cationic AMPs is an important aspect of their function remains underexplored.

Since microbial membranes are the primary targets of AMPs, the prerequisite for their action is the ability to reach the outer membrane of Gram-negative bacteria to cause its permeabilization and cell lysis (2, 68). It is widely held that bacterial CPS, including that of *K. pneumoniae*, can restrict AMP access to the bacterial membrane. In this regard, it has been shown that an acapsular mutant of *K. pneumoniae* was more sensitive to defensins than the wild-type strain (71). Furthermore, free exopolysaccharide, which can be released from the bacterial surface, binds AMPs, neutralizing their bactericidal effect (72), suggesting that trapping AMPs by anionic CPS is a general mechanism by which CPS counteracts AMPs. Thus, both complement proteins and AMPs have a limited ability to reach and disrupt the outer bacterial membrane, indicating that through the CPS, *K. pneumoniae* has developed a strategy to evade the lethality of two main host defenses systems. One important property of rdPF4, which otherwise closely resembles AMPs in traits such as small size and abundance of positively charged and hydrophobic residues, is the lack of cytotoxicity for host cells, which is common among many AMPs. We previously demonstrated that, despite its beneficial direct bactericidal effects and phagocytosis-promoting activity, LL-37 can damage the host cell membrane, leading to cytoplasmic leakage, whereas rdPF4 is not cytotoxic even at very high concentrations (28). In addition to binding to CPS, rdPF4 also binds to the negatively charged bacterial wall of nonencapsulated bacteria; however, it does not permeabilize bacterial membranes but instead acts as an opsonin (28, 30). The reason why rdPF4 does not damage bacteria and host cell membranes remains speculative. The C-terminal region of rdPF4 spanning residues 57 through 70 and containing the CR3-binding site adopts an amphipathic α-helical conformation, typical of many AMPs (73), (68). However, even though the synthetic peptide duplicating this sequence exhibits bactericidal effect (74), the entire molecule lacks this activity. It is well established that both positively charged and hydrophobic residues, which are on opposite sides of the amphipathic α-helix, are essential for their interaction with and insertion into bacterial membranes (73, 75). In the PF4 monomer (PDB ID: 1F9Q), the C-terminal α-helix is not free but embedded in the protein scaffold, exposing Lys61, Lys62, and Lys65 on the molecule’s surface while burying several hydrophobic residues (Ile63, Ile64, and Leu67) within the interface between the helix and the rest of the protein. This arrangement may prevent these residues from reaching the membrane, thereby avoiding disruption of the lipid bilayer. The combination of other features, including charge, hydrophobicity, peptide length, and the conformation of rdPF4, can enhance its safety for host cells while increasing affinity for CR3.

As the main virulence factor, the capsule has been targeted for the treatment of severe bacterial infections, including *K. pneumoniae* CPS (reviewed in (5, 47, 76)). Among strategies to overcome CPS resistance to phagocytosis, anticapsular antibodies, phage-derived capsule depolymerases, and small-molecule inhibitors of capsule biosynthesis have been investigated for treating bacterial infections, particularly those caused by drug-resistant pathogens. In addition to these approaches, targeting the capsule of *K. pneumoniae* with rdPF4 could potentially be developed into an effective therapy.

## Acknowledgments

This work was supported by a grant from the Flinn Foundation and partly by an NIH grant HL63199. We acknowledge the use of instruments within the Biosciences Advanced Light Microscopy Facility at Arizona State University.

## Author contribution

Conceived and designed the analysis: **N.P**., **T.U.**

Collecting data and analysis: **N.P., I.K., J.A., C.D., A.B., D.R.**

Funding acquisition: **T.U., N.P.**

Writing, Review, and Editing the paper: **T.U.**, **N.P.**

