## Supplemental Material for "TARGETING THE CAPSULE OF *KLEBSIELLA PNEUMONIAE* WITH A CATIONIC CR3-BINDING PROTEIN ENHANCES PHAGOCYTOSIS AND PROMOTES BACTERIAL CLEARANCE AND SURVIVAL IN A MOUSE SEPSIS MODEL"

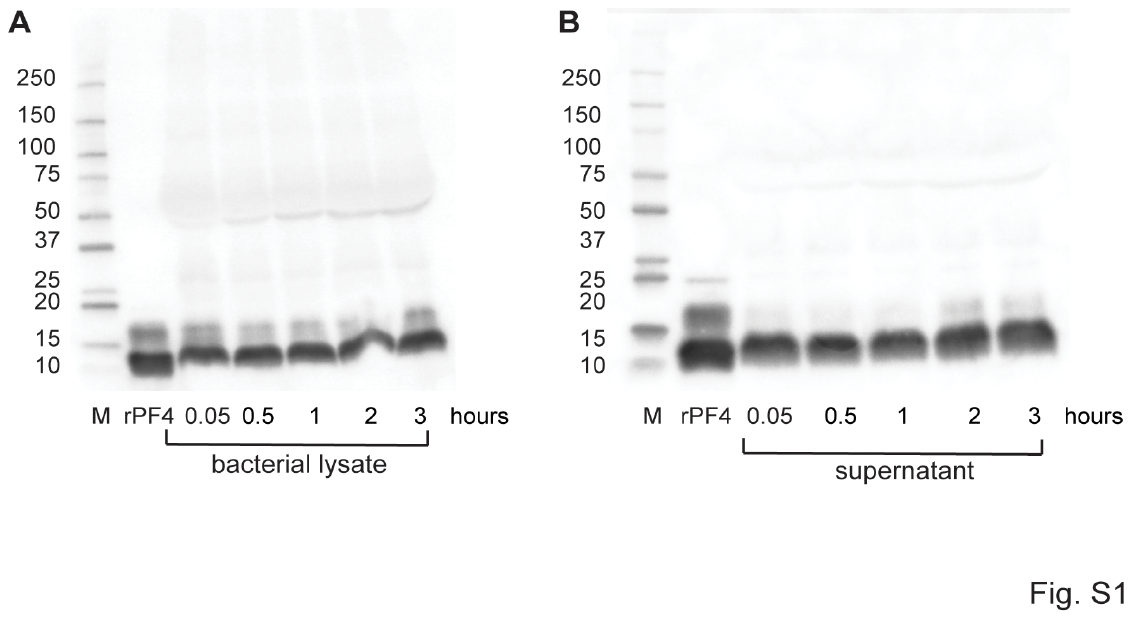

**Figure S1.** rdPF4 stability in human plasma. Overnight culture of *K. pneumoniae* was diluted 1:100 in human plasma. Bacterial suspensions (1 ml) were incubated with 100 µg/ml rdPF4 for 0.5-3 h at 37 ºC. Bacterial cells (**A**) and supernatants (**B**) were analyzed for the presence of rPF4 by Western blotting using a rabbit polyclonal anti-PF4 antibody and a goat anti-rabbit secondary antibody. M, protein standards, rPF4, purified recombinant dimeric PF4.

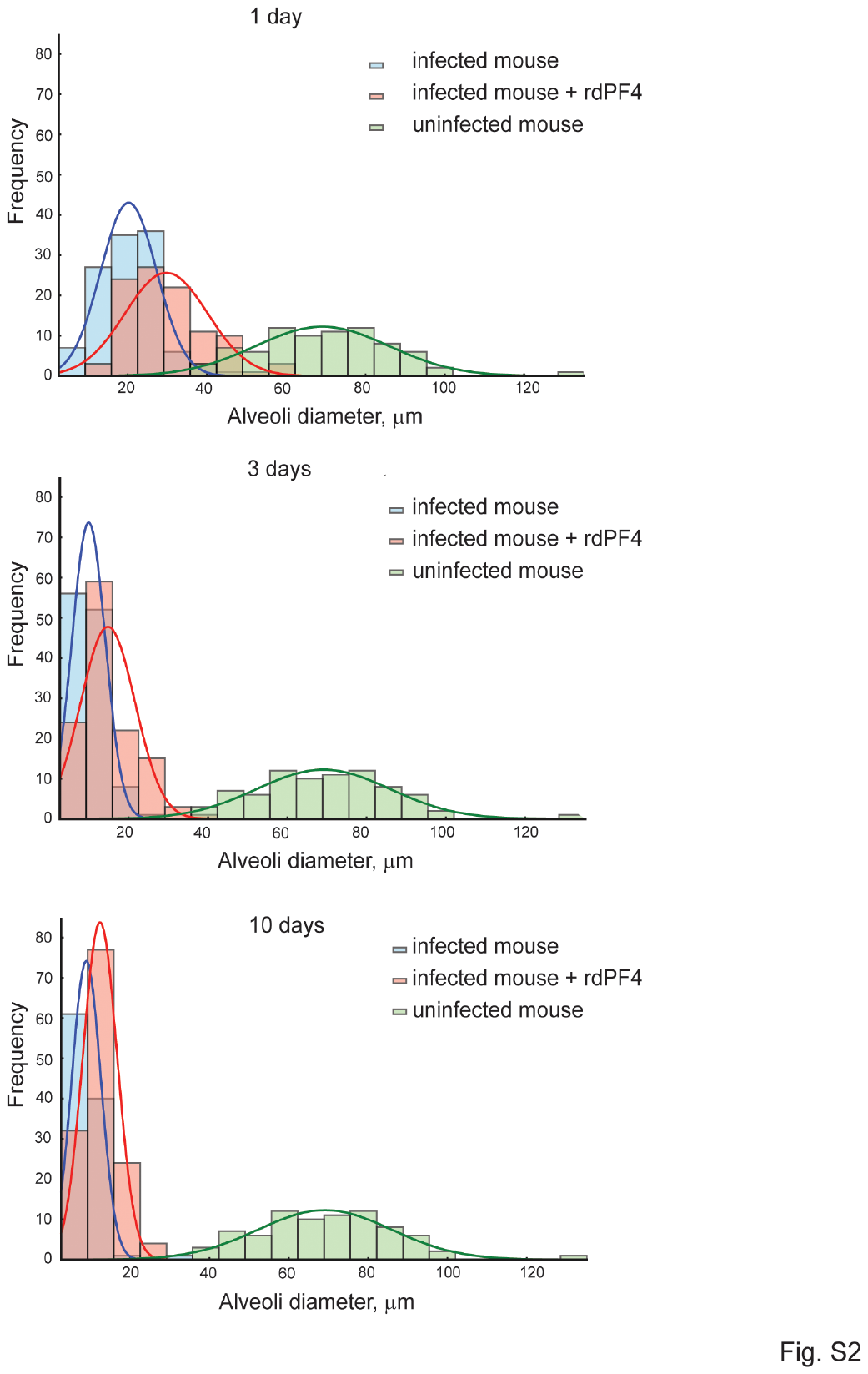

**Figure S2.** Frequency distribution of the total alveoli diameter in lungs of uninfected, infected, and infected mice treated with rdPF4 1, 3, and 10 days after infection with low-virulence *K. pneumoniae*.

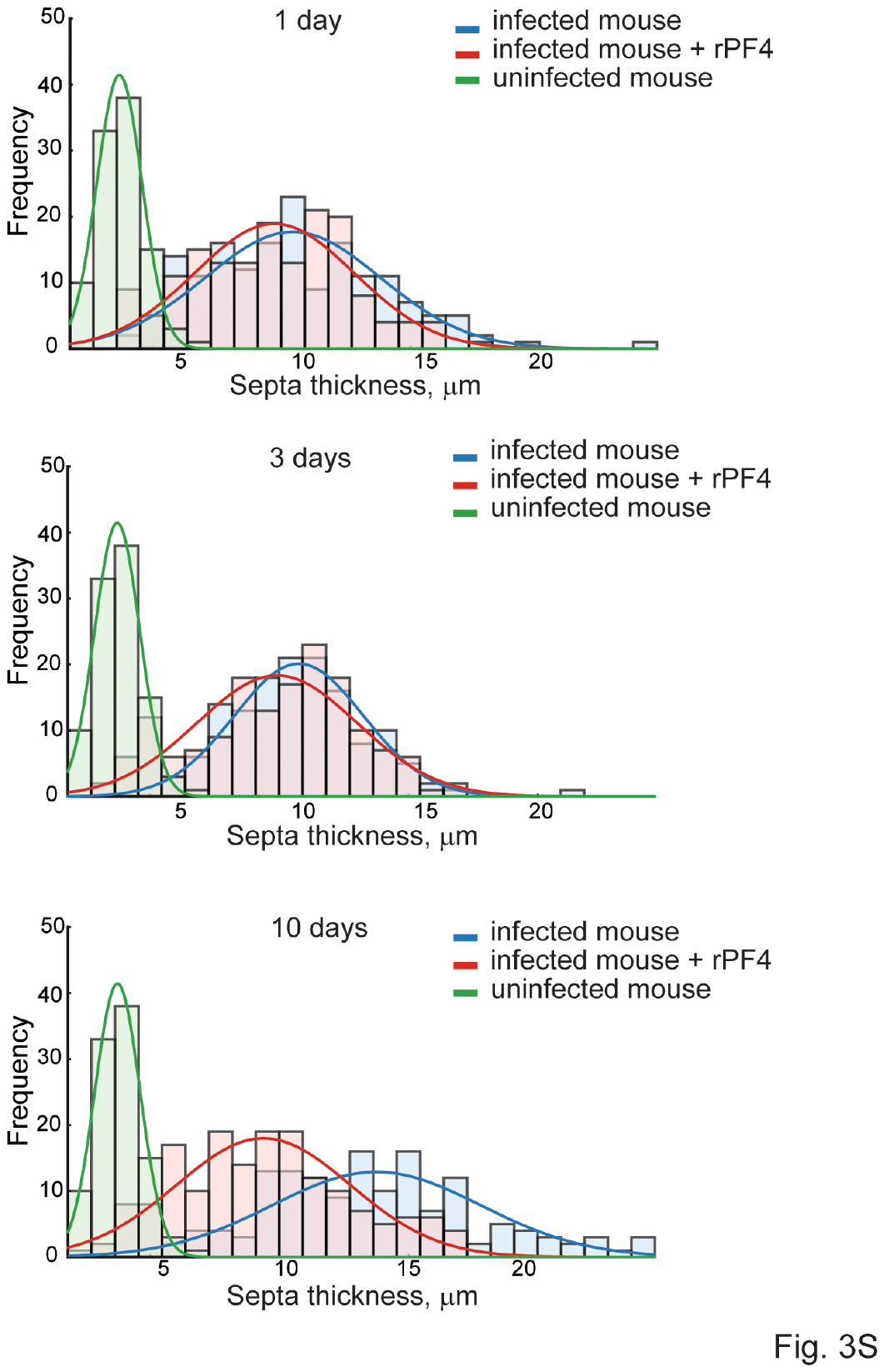

**Figure S3.** Frequency distribution of the septa thickness in lungs of uninfected, infected, and infected mice treated with rdPF4 1, 3, and 10 days after infection with low-virulence *K. pneumoniae*.

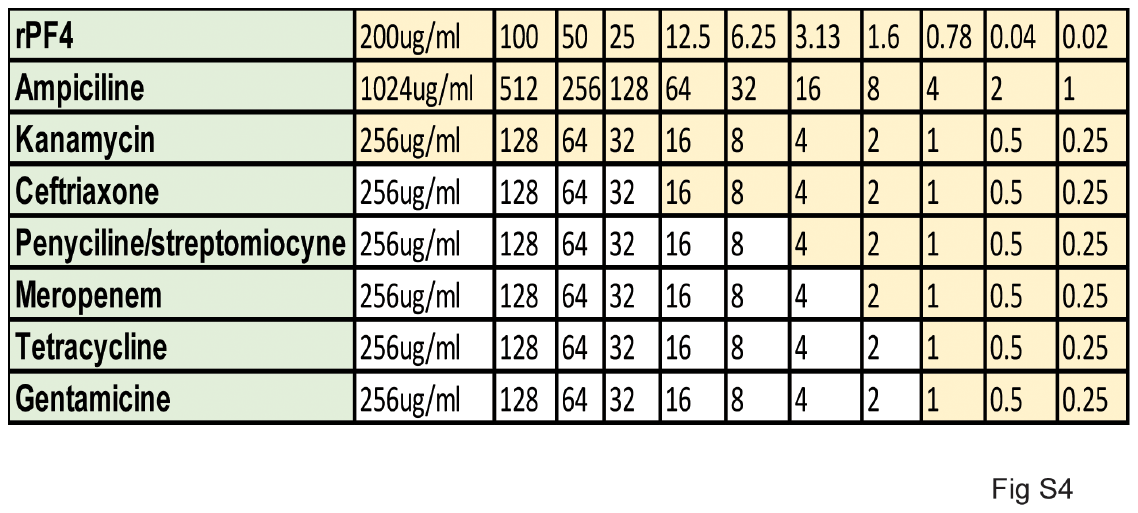

**Figure S4**. Minimum Inhibitory Concentration (MIC) of 7 antibiotics and rdPF4 against the BAA-1705 strain.

**Supplemental Table**

Counts of bacteria in the peritoneum, lungs, liver, and blood of mice 24 hours after infecting neutrophil-depleted mice with 3x10^7^ CFU and macrophage-depleted mice with 1x10^7^ CFU of lv*Kp*.

|  | **Nondepleted, CFU** (n=4) | **Neutrophil-depleted, CFU** (n=4) | **Fold, CFU increase in neutrophil-depleted vs nondepleted mice** |
| --- | --- | --- | --- |
| Peritoneal lavage | 4.6x10^7^ ± 3.2x10^7^ | 3.44x10^9^ ± 6.8x10^8^ | 75 ± 21 |
| Lung | 2.6x10^5^ ± 1.7x10^5^ | 2.84x10^7^ ± 1.1x10^7^ | 109 ± 64 |
| Liver | 2.5x10^6^ ± 1.4x10^6^ | 6.8x10^7^ ± 4.5x10^6^ | 27 ± 3 |
| Blood | 2.4x10^5^ ± 1.5x10^5^ | 5.04x10^8^ ± 3.4x10^8^ | 2136 ±2266 |

|  | **Nondepleted, CFU** (n=4-7) | **Macrophage-depleted, CFU** (n=6) | **Fold, CFU increase in macrophage-depleted vs nondepleted mice** |
| --- | --- | --- | --- |
| Peritoneal lavage | 2.6x10^4^ ± 4.1x10^4^ | 2.8x10^8^± 4.8x10^8^ | 10910 ±11707 |
| Lung | 4.4x10^2^ ± 6.0x10^2^ | 1.6x10^3^± 3.2x10^3^ | 3.6 ±5.3 |
| Liver | 1.3x10^5^ ± 2.4x10^5^ | 2.8x10^8^± 3.9x10^8^ | 2121 ± 1625 |
| Blood | 1.1x10^3^ ± 1.3x10^3^ | 2.0x10^6^±4.2x10^6^ | 1870 ± 3230 |
